# Modeling second-order boundary perception: A machine learning approach

**DOI:** 10.1101/369041

**Authors:** Christopher DiMattina, Curtis L. Baker

**Affiliations:** Computational Perception Laboratory, Department of Psychology, Florida Gulf Coast University, Fort Myers, FL, USA 33965-6565; McGill Vision Research Unit, Department of Ophthalmology, McGill University, Montreal, QC, Canada H3G1A4

## Abstract

**Background:** Visual pattern detection and discrimination are essential first steps for scene analysis. Numerous human psychophysical studies have modeled visual pattern detection and discrimination by estimating linear templates for classifying noisy stimuli defined by spatial variations in pixel intensities. However, such methods are poorly suited to understanding sensory processing mechanisms for complex visual stimuli such as second-order boundaries defined by spatial differences in contrast or texture.

**Methodology / Principal Findings:** We introduce a novel machine learning framework for modeling human perception of second-order visual stimuli, using image-computable hierarchical neural network models fit directly to psychophysical trial data. This framework is applied to modeling visual processing of boundaries defined by differences in the contrast of a carrier texture pattern, in two different psychophysical tasks: (1) boundary orientation identification, and (2) fine orientation discrimination. Cross-validation analysis is employed to optimize model hyper-parameters, and demonstrate that these models are able to accurately predict human performance on novel stimulus sets not used for fitting model parameters. We find that, like the ideal observer, human observers take a region-based approach to the orientation identification task, while taking an edge-based approach to the fine orientation discrimination task. How observers integrate contrast modulation across orientation channels is investigated by fitting psychophysical data with two models representing competing hypotheses, revealing a preference for a model which combines multiple orientations at the earliest possible stage. Our results suggest that this machine learning approach has much potential to advance the study of second-order visual processing, and we outline future steps towards generalizing the method to modeling visual segmentation of natural texture boundaries.

**Conclusions / Significance:** This study demonstrates how machine learning methodology can be fruitfully applied to psychophysical studies of second-order visual processing.

**Author Summary:** Many naturally occurring visual boundaries are defined by spatial differences in features other than luminance, for example by differences in texture or contrast. Quantitative models of such “second-order” boundary perception cannot be estimated using the standard regression techniques (known as “classification images”) commonly applied to “first-order”, luminance-defined stimuli. Here we present a novel machine learning approach to modeling second-order boundary perception using hierarchical neural networks. In contrast to previous quantitative studies of second-order boundary perception, we directly estimate network model parameters using psychophysical trial data. We demonstrate that our method can reveal different spatial summation strategies that human observers utilize for different kinds of second-order boundary perception tasks, and can be used to compare competing hypotheses of how contrast modulation is integrated across orientation channels. We outline extensions of the methodology to other kinds of second-order boundaries, including those in natural images.

## Introduction

Many of the most common functions of sensory systems involve detection, identification or discrimination of particular stimuli. For example, a critically important ability of early visual processing is to segment images into distinct regions, which may correspond to cohesive surfaces or objects (Marr, 1982). To better understand the underlying mechanisms, it would be useful to fit biologically plausible models of surface segmentation to psychophysical data, and assess how well such models can account for independent datasets.

A widely used approach to modeling human psychophysics has been to assume that performance can be understood in terms of how well a visual stimulus matches an internal “template”, typically modeled as a linear spatial filter. This approach uses high-dimensional regression models fit to data from tasks in which a subject detects or classifies a sensory stimulus presented with superimposed white noise (Ahumada, 1996, 2002; Neri & Levi, 2006; Murray, 2011), to recover an estimate of the linear filter that best accounts for the data. This system-identification (SI) approach to psychophysics, often called “psychophysical reverse correlation” or “classification image analysis”, parallels neurophysiology studies that fit linear filter models to neural responses elicited by white noise stimuli (e.g. DeAngelis, Ohzawa, & Freeman 1993; Chichilnisky, 2001). Psychophysical SI methods have been applied in varied problem domains and yielded valuable insight into perceptual mechanisms (Abbey & Eckstein, 2002; Eckstein, Shimozaki, & Abbey, 2002; Li, Levi, & Klein, 2004; Neri & Levi, 2006; Morgenstern & Elder, 2012; McIlhagga & May, 2012; Kurki, Saarinen, & Hyvarinen, 2014).

One limitation of most previous psychophysical system identification studies has been the use of stimuli that are typically defined by spatial or temporal variations in first-order statistics like luminance or color. However, many natural visual stimuli such as occlusion boundaries (Fig. 1a, top) are defined by spatial variations in higher-order image statistics, e.g. texture or contrast, rather than simple luminance differences (Schofield, 2000; Konishi et al., 2003; Johnson & Baker, 2004; Martin, Fowlkes, & Malik, 2004; DiMattina, Fox, & Lewicki, 2012). Such “second-order” boundaries (Fig. 1a, bottom) have been of particular interest because their detection cannot be explained by simple linear filters (Baker, 1999; Landy & Graham, 2004), and consequently human performance cannot be readily modeled using conventional psychophysical SI methods. More generally, there is a need for a psychophysical modeling approach that can handle a broader range of stimuli and tasks.

**Figure 1:**
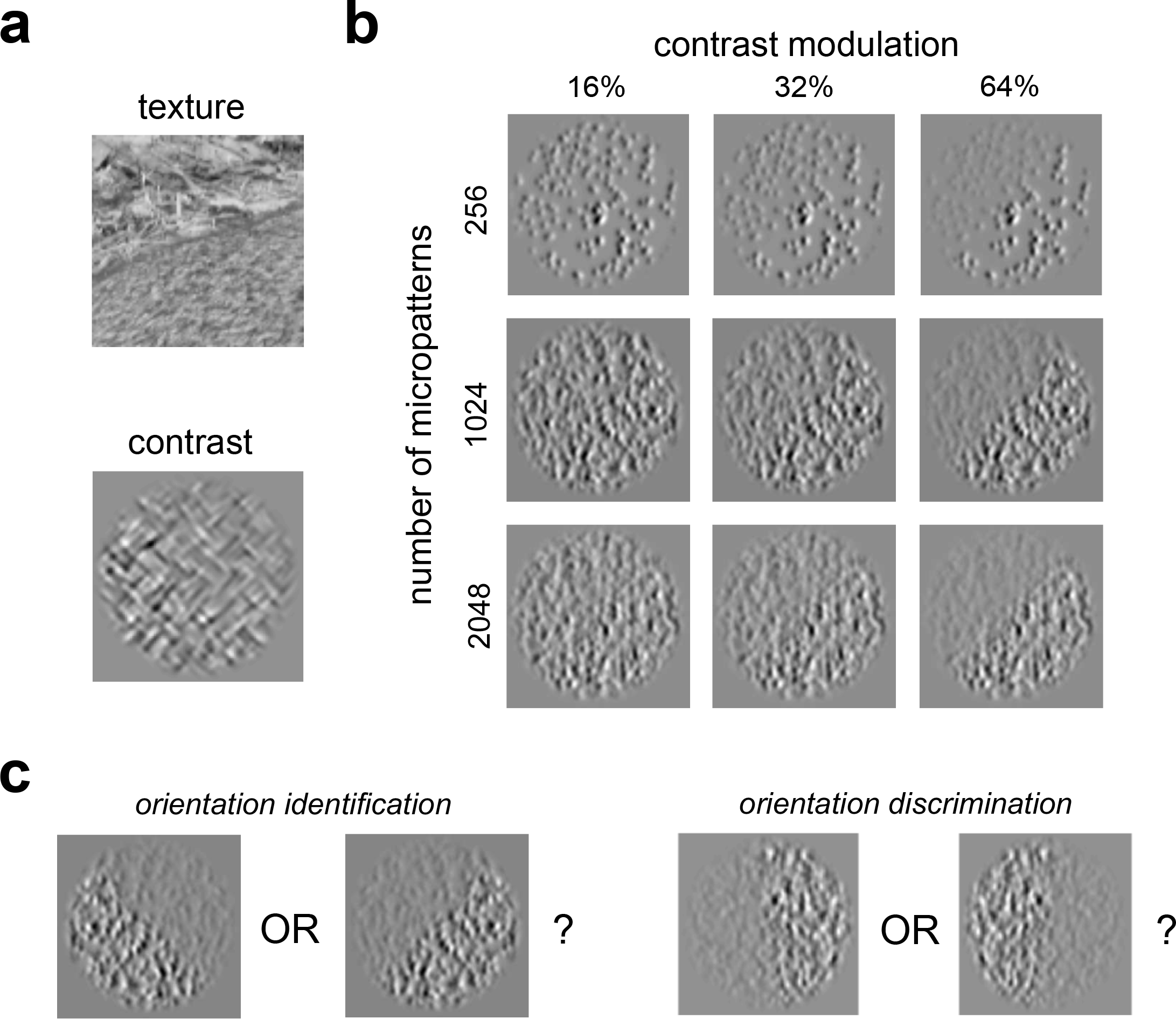
Examples of second-order boundary stimuli, with varying texture density and modulation depth. **(a)** *Top:* A natural occlusion boundary having similar average luminance on both sides, but clearly visible differences in texture. *Bottom:* A contrast-defined boundary with identical mean luminance in each region, but different contrasts of the texture elements. In this example the contrast modulation envelope has a left-oblique (−45 deg. w.r.t vertical) orientation. **(b)** Examples of contrast-modulated micropattern stimuli, for three densities of micropatterns and three modulation depths, all having a right-oblique (+45 deg.) boundary orientation. **(c)** Schematic illustration of the two psychophysical tasks used here (*orientation identification*: Experiments 1 and 3, *orientation discrimination*: Experiment 2), in both cases two-alternative forced-choice judgements of left-vs. right-oblique boundary orientation.

Here we present a novel machine learning method to extend psychophysical SI to more complex naturalistic stimuli such as second-order boundaries. Our general approach is to fit image-computable hierarchical neural network models of second-order vision (Fig. 2) directly to psychophysical trial data, similar to recent studies using multi-stage models to characterize nonlinear responses in sensory neurons (Willmore, Prenger, & Gallant, 2010; Vintch, Movshon, & Simoncelli, 2015; Harper et al. 2016; Rowenkamp & Sharpee, 2017). Our application of this framework generalizes a class of biologically plausible “Filter-Rectify-Filter” (FRF) models that have often been invoked to explan second-order processing (Landy & Graham, 2004). However, in contrast to previous general schemes for second-order vision (e.g. Malik & Perona, 1990; Bergen & Landy, 1991; Graham, Beck & Sutter, 1992) and quantitative models (Zavitz & Baker, 2013; Westrick & Landy, 2013), we take a machine learning approach in which we estimate the defining model parameters by direct fitting to psychophysical trial data. This permits more incisive tests of predictions on novel datasets in order to validate our models, and to evaluate competing models.

**Figure 2:**
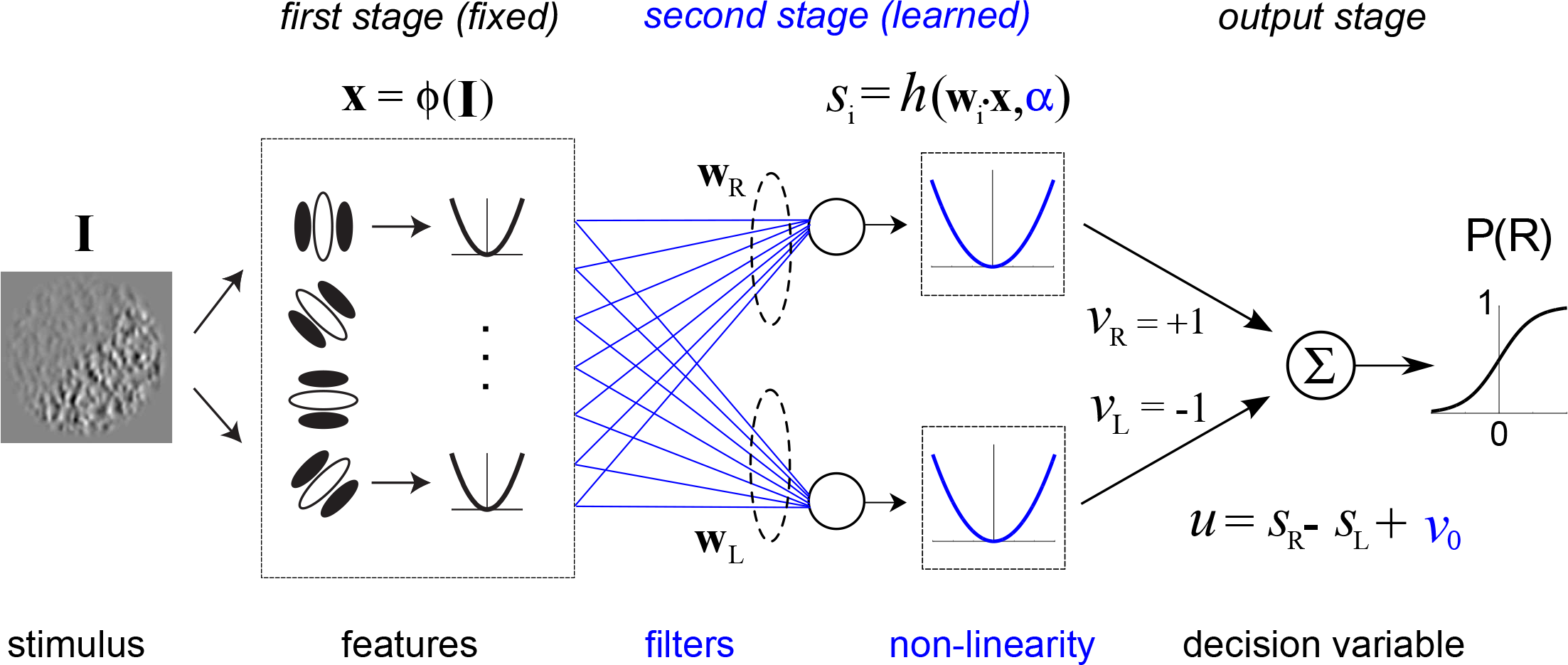
Neural network model implementing a FRF (Filter-Rectify-Filter) arrangement for texture segmentation. Parameters shown in blue are learned from data, those in black are fixed. Stimulus image **I** is filtered by a bank of first-stage energy filters, resembling V1 complex cells, whose downsampled responses provide a feature vector input **x** to two second-stage filters. An optimization algorithm adjusts connection weights **w**_L_, **w**_R_ (blue lines), producing second-stage filters which are selective for left-oblique (L) or right-oblique (R) contrast-defined boundaries, to give responses consistent with human psychophysical data. The output of each second-stage filter is passed through a nonlinearity *h*(*u*) = |*u*|^*α*^ (blue curve) whose shape parameter α is also estimated from the psychophysical data. Fixed output weights *v*_R_ = +1, *v*_L_ = −1 lead to the decision variable *u* = *s*_R_ − *s*_L_ + *v*_0_, which is input to a sigmoid function to determine the probability of the observer classifying a given boundary as being right-oblique.

We focus on contrast-defined second-order boundaries (Fig. 1a, b) since they are well studied psychophysically (Dakin & Mareschal, 2000; Schofield & Georgeson, 1999, 2003; Ledgeway & Smith, 1994; Ellemberg, Allen, & Hess 2006), and there are neurons in the early mammalian visual cortex that give selective responses to contrast modulation stimuli (Mareschal & Baker, 1998; Tanaka & Ohzawa, 2006; Rosenberg et al, 2010; Li et al., 2014). We model performance on two tasks making use of contrast-defined boundaries: (1) identification of boundary orientation, and (2) boundary orientation discrimination (Fig. 1c). We find that subjects utilize spatial information in markedly different ways for these two tasks, employing “region-based” processing for the orientation identification task and “edge-based” processing for the orientation discrimination task, in both cases consistent with an ideal observer.

We further demonstrate the power of this methodology for evaluating competing hypotheses regarding the nature of channel summation within the FRF scheme (Kingdom, Prins, & Hayes, 2003; Motoyoshi & Nishida, 2004; Motoyoshi & Kingdom, 2007; Westrick & Landy 2013) by comparing the ability of two different model architectures to fit human performance on a contrast edge detection task in which the carrier pattern contains elements with two orientations. Post-hoc Bayesian model comparison (Schwarz, 1978; Kass & Raferty, 1995; Bishop, 2006) demonstrates a preference for a model in which carrier pattern orientation is integrated across channels by the second-stage filters, consistent with previous psychophysical studies (e.g., Motoyoshi & Nishida, 2004).

Finally, we discuss potential applications of this methodology to future quantitative studies of first- and second-order cue integration (Allard & Faubert, 2007; Saarela & Landy, 2012; Hutchinson, Ledgeway, & Baker, 2016) and natural boundary detection (McDermott, 2004; DiMattina et al., 2012). We suggest that the application of machine learning methodology similar to that presented here can potentially help to better quantify visual processing of natural occlusion boundaries and complex natural visual stimuli more generally.

## Results

Observers performed two different psychophysical tasks making use of contrast-modulated boundary stimuli like those shown in Fig. 1 (see Methods). In the first task, orientation-identification (Fig. 1c, left), observers determined in which of two oblique orientations (−/+ 45 deg, L/R) a near-threshold contrast boundary was presented, with varying amounts of contrast modulation (Experiments 1 and 3). We also employed an orientation-discrimination task (Fig. 1c, right) in which observers indicated the orientation (L/R) of a supra-threshold contrast boundary, whose orientation was perturbed slightly from vertical by varying amounts (−/+ 6-7 deg.) (Experiment 2). Trial-wise responses obtained from these experiments were used to fit and evaluate a model (Fig. 2) implementing a “Filter-Rectifty-Filter” (FRF) operation (Landy & Graham, 2004) on each stimulus image, with the goal of characterizing the nature of spatial summation for perception of contrast boundaries, as well as the shape of the second-stage filter nonlinearity. We describe the model in greater depth before presenting the experimental results.

### Modeling framework

The model used to fit psychophysical trial data (Fig. 2) employed a first stage comprised of a bank of fine-scale Gabor filters that compute local oriented energy in a manner similar to V1 complex cells (Adelson & Bergen, 1985). The second stage consisted of a pair of filters, representing the two task alternatives (L/R), each instantiated by a weighted sum of the normalized outputs of the first-stage filters. In contrast to the first-stage filters, these second-stage filters analyze the image at a more coarse spatial scale, integrating variations in contrast across the boundary. Finally, the outputs of these two second-stage filters are passed through a static non-linearity *h*(*u*) = |u|^*α*^ and then compared to decide on a trial-by-trial basis if a given stimulus boundary appeared more likely to be tilted left or right (L/R), i.e. the probability 0 < P(R) < 1 of the stimulus being classified by the observer as R. The main goal of Experiments 1 and 2 was to estimate the shapes of the second-stage filters.

To reduce model parameter space dimensionality, we performed downsampling between the first- and second-stage filters, i.e. pooling of spatially adjacent first-stage filter responses. Here we implemented three alternative methods: (1) simple averaging (AVG), (2) maximum (MAX) (Goodfellow, Bengio, & Courville, 2016), and (3) sub-sampling (SUB) (Bergen & Landy, 1991). In Experiment 1 we considered all three pooling rules; most analyses utilized 16×16 downsampling, but 3 downsampling sizes (22×22, 16×16 and 8×8) were compared. In Experiments 2 and 3 we considered only AVG pooling and a single downsampling size (Experiment 2: 22×22, Experiment 3: 12×12).

We implement the model’s final (L/R) classification in two alternative ways: In the *deterministic* model (DET), if the output of the R filter is larger than that of the L filter (plus the bias term) so that P(R) > 0.5, the stimulus will be classified as R, otherwise it is classified as L. In the *stochastic* model (STO), the output P(R) specifies only the probability of classifying the stimulus as R or L, thus introducing randomness to the model without requiring an additional free parameter.

A common approach to make the estimation problem more tractable is to utilize reasonable prior constraints on the model parameter space (Mineault, Barthelme, & Pack, 2009; Wu, David & Gallant, 2006; Park & Pillow, 2011). Here we use two kinds of Bayesian priors: (1) ridge-regression (Hoerl & Kennard, 1970; Bishop 2006) to penalize large weights (*ridge*), and (2) ridge-regression combined with an additional smoothness prior (Mineault et al., 2009) to penalize excessively “rough” second-stage filter weight profiles (*ridge* + *smooth*). Additional details of the model are given in Methods.

### Experiment 1: Boundary orientation identification

#### Stimuli and task

Observers viewed contrast-defined texture boundaries with varying degrees of modulation depth (Fig. 1b), and performed forced-choice judgments of whether the boundary was oriented right- or left-oblique (45 degrees clockwise or counterclockwise, Fig.1c left). In Experiment 1-VAR (7 observers), a method of constant stimuli was used to vary contrast modulation depth over a range spanning values near threshold but also including some levels producing near-perfect, or chance, performance. In another version (Experiment 1-FIX, 3 observers) the contrast modulation depth was held constant at a near-threshold level.

#### Model fit to representative observers

Fig. 3a shows the results of fitting the model (16×16 downsampling, AVG pooling) to data from one representative observer (AMA) in Experiment 1-VAR. The top panels show estimated second-stage filter weights as image maps across the spatial extent of the stimuli, with pixel color indicating weight values. The black dashed lines in the bottom plots show mean 1-D profiles, obtained by collapsing the 2-D weight maps orthogonally to the boundary orientations, whereas thick solid lines (red = *ridge*, green = *ridge* + *smooth*) show profiles obtained from over 30 bootstrap-resampled training sets. We see that these mean 1-D profiles are nearly indistinguishable from one another, all resembling a step-like function (see below). While many of the individual 2-D weights can be quite variable (Supplementary Fig. S7), the 1-D weight profile magnitudes lie well above zero (Fig. 3, thin dashed lines show +/- 1 SEM). The addition of the smoothing prior (*ridge* + *smooth*) results in weighting functions that are very similar, but less “noisy” in appearance, consistent with previous results with classification images and receptive field modeling (Mineault et al., 2009; Park and Pillow, 2011). Furthermore, all three pooling rules (AVG, MAX, SUB) yielded consistent 2-D and 1-D weight profiles (Supplementary Fig. S8).

**Figure 3:**
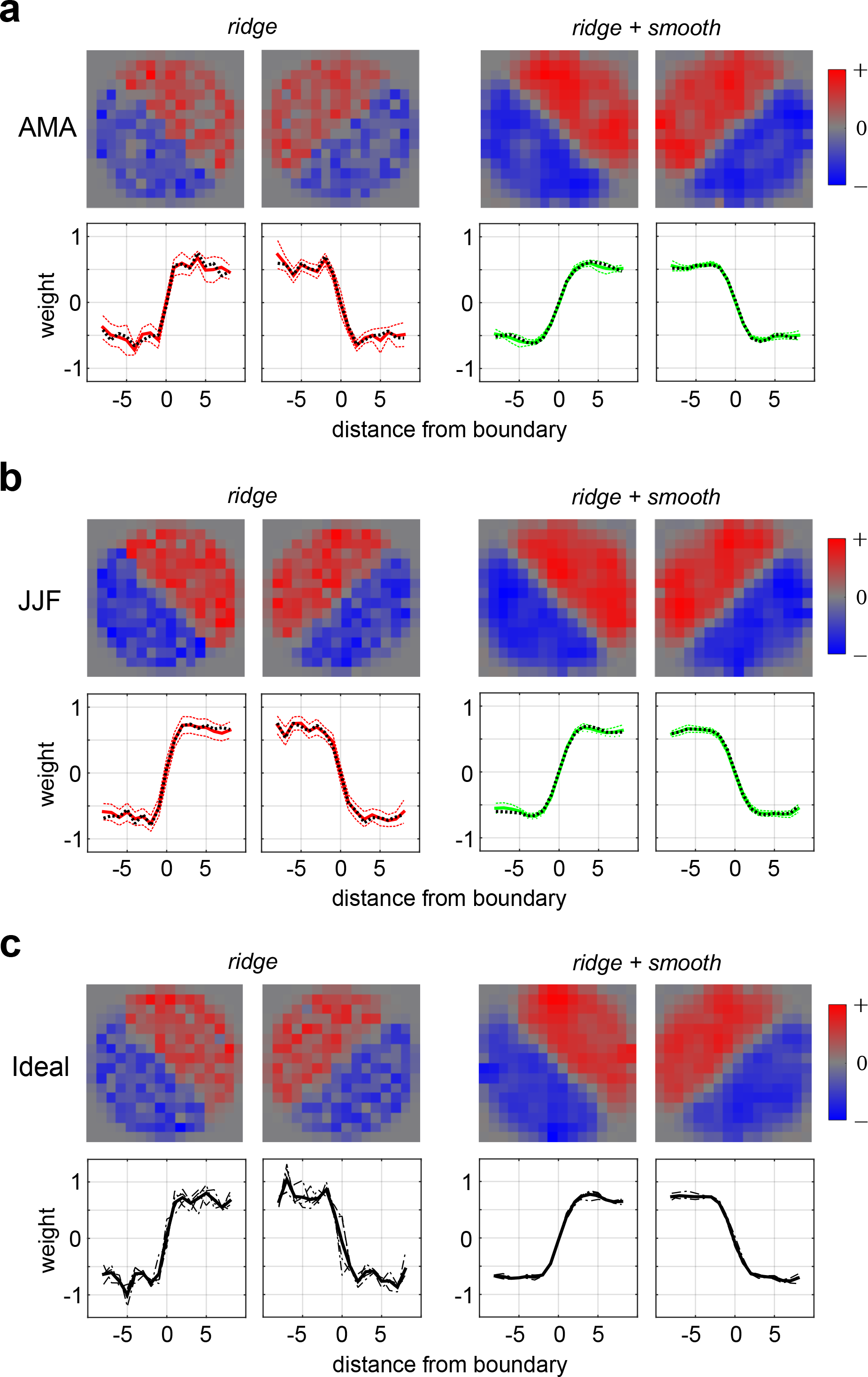
Estimated second-stage filter weights for two representative observers in Experiment 1 (orientation identification). **(a)** Second-stage filter weights learned by the model for representative observer AMA in Experiment 1-VAR (varying modulation depth) for two different priors (left: *ridge*; right: *ridge + smooth*) with 16×16 AVG downsampling. Top panels show the 2-D filter weights (averaged over 4 training folds) and bottom panels show these 2-D weights collapsed into 1-D profiles (black dashed lines) by averaging along the matrix diagonals (left-oblique) or anti-diagonals (right-oblique). Thick lines (red: *ridge*; green: *ridge + smooth*) denote averages over 30 resampled bootstrapped training sets, and thin dashed lines show +/- 1 SEM. **(b)** Same as (a) but for observer JJF in Experiment 1-FIX (fixed, near-threshold modulation depth). **(c)** Results for ideal observer for Experiment 1-VAR. Organization as in (a), (b) except thick black lines denote averages over 4 training folds and thin dashed lines show fits to individual folds.

Fig. 3b shows the model fit to data from a representative observer (JJF) in Experiment 1-FIX. In this version of the experiment, the contrast modulation depth was kept constant at approximately threshold level for each observer (Supplementary Table S1), similar to most previous work in the classification image literature (Abbey & Eckstein, 2002; Murray, 2011). Here we see very similar results as for the observer in Experiment 1-VAR, also showing a step-like function for this task. As before, this result was robust to the pooling rule (Supplementary Fig. S9).

To assess the extent to which observers were employing the same spatial integration as an ideal observer, we fit the model to stimulus category labels rather than observer responses. The 2-D weight maps and 1-D profiles obtained for an ideal observer in Experiment 1-VAR (Fig. 3c), reveal that the ideal weight map is a step-like function like that seen in the human results. Similar results for an ideal observer fit to stimulus category labels in Experiment 1-FIX are shown in Supplementary Fig. S10.

#### Edge-based vs. region-based analysis

When finding a boundary between adjacent textures, the visual system might make use only of texture elements close to the boundary (“edge-based”), or it might utilize the entirety of both adjacent texture regions (“region-based”). A number of previous studies have investigated this question for textures defined by orientation differences (Nothdurft, 1985; Gurnsey & Laundry, 1992; Wolfson & Landy, 1998), but to the best of our knowledge this issue has not been addressed systematically for contrast-defined textures. The results from two individual observers (Fig. 3a, b) as well as an ideal observer (Fig. 3c) indicate that the entire texture regions on either side of the boundary are contributing to performance, giving step-like 1-D profiles - all indicative of “region-based” processing.

To examine the robustness of this result across observers and for different model variants, Fig. 4 shows the 1-D averaged profiles of second-stage filter weights for all observers (thin colored lines) and group averages (thick black lines) in Experiment 1-VAR (Fig. 4a) and Experiment 1-FIX (Fig. 4b). We see that the magnitudes of the 1-D weights are approximately constant as one moves from the diagonal to the boundary, indicative of region-based processing, and that these results are also consistent for all three pooling rules (AVG, MAX, SUB). To explore whether the step-like 1-D profile result is consistent with an ideal observer, plots of the normalized (scaled to unit norm) ideal observer 1-D weight profiles (30 bootstraps) are shown in Fig. 5a as thick black lines, with 95% confidence intervals (+/- 1.96 SEM) plotted as dashed black lines. We see that the normalized 1-D weight profiles for individual observers (dashed color lines), and for means over observers (thick red dashed lines) generally lie within the 95% confidence intervals of the ideal observer. Therefore, the performance of the human observers is very much consistent with that of the ideal observer, both of which give similar step-like profiles and thus seem to integrate contrast information across the whole region.

**Figure 4:**
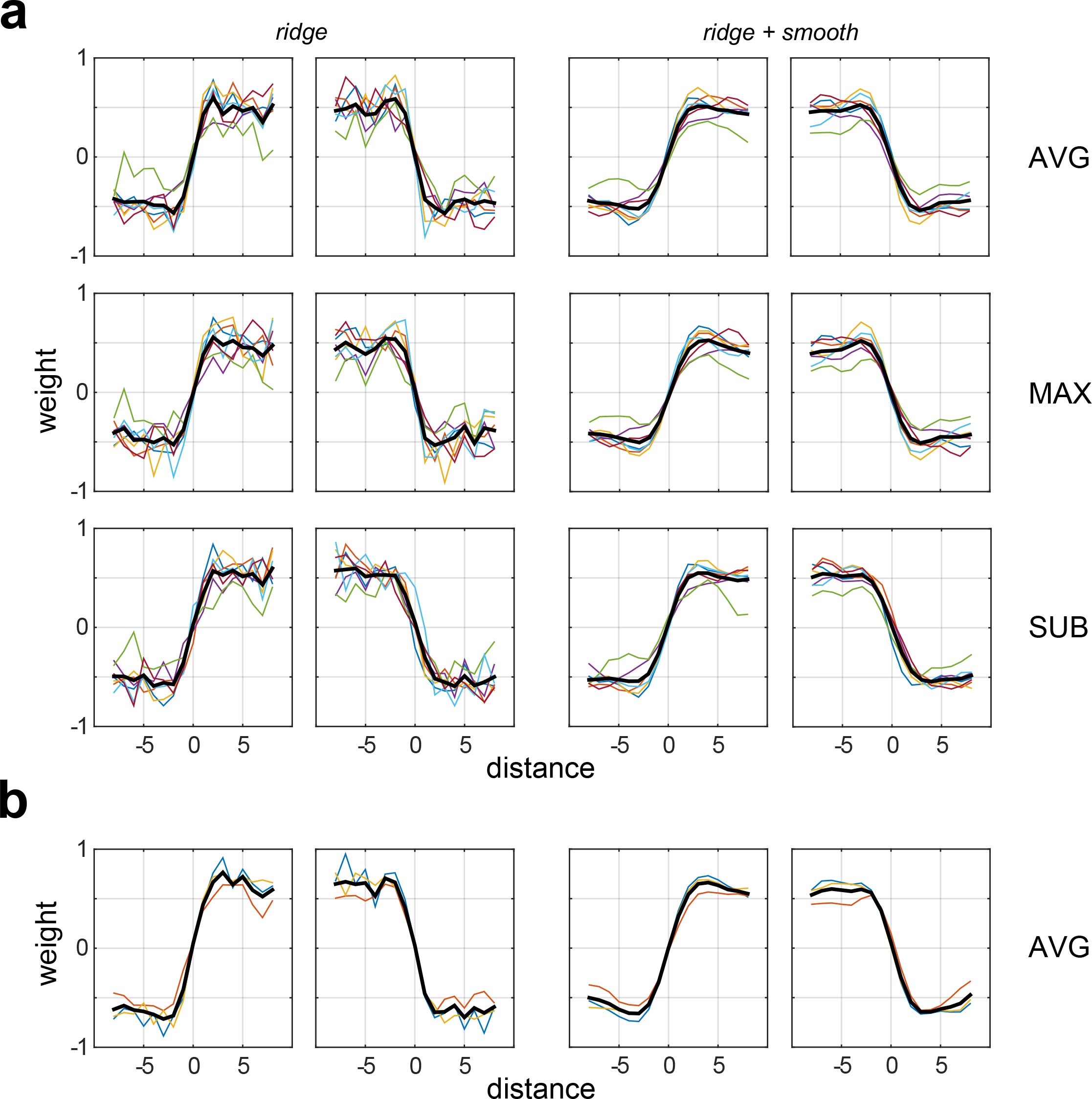
1-D profiles of the second-stage filter weights for all observers in Experiment 1. **(a)** 1-D profiles for individual observers in Experiment1-VAR (colored lines) plotted with the average across observers (thick black lines) for three sampling rules (AVG, MAX, SUB) and both priors. **(b)** Same as (a) but for Experiment 1-FIX.

To test whether our finding of region-based processing was robust to increasing the carrier spatial frequency, we ran an additional control version of the experiment on 3 naïve observers (Experiment 1-HFC) in which we reduced the micropattern size from 32 to 8 pixels (while increasing the density by a factor of 4). Such a stimulus would potentially engage many more contrast-modulation tuned neurons having receptive fields localized to the boundary (Li et al., 2014), potentially revealing edge-based processing. However, even with the high-frequency carrier we still observed region-based processing (Fig. 5b), consistent with an ideal observer.

**Figure 5:**
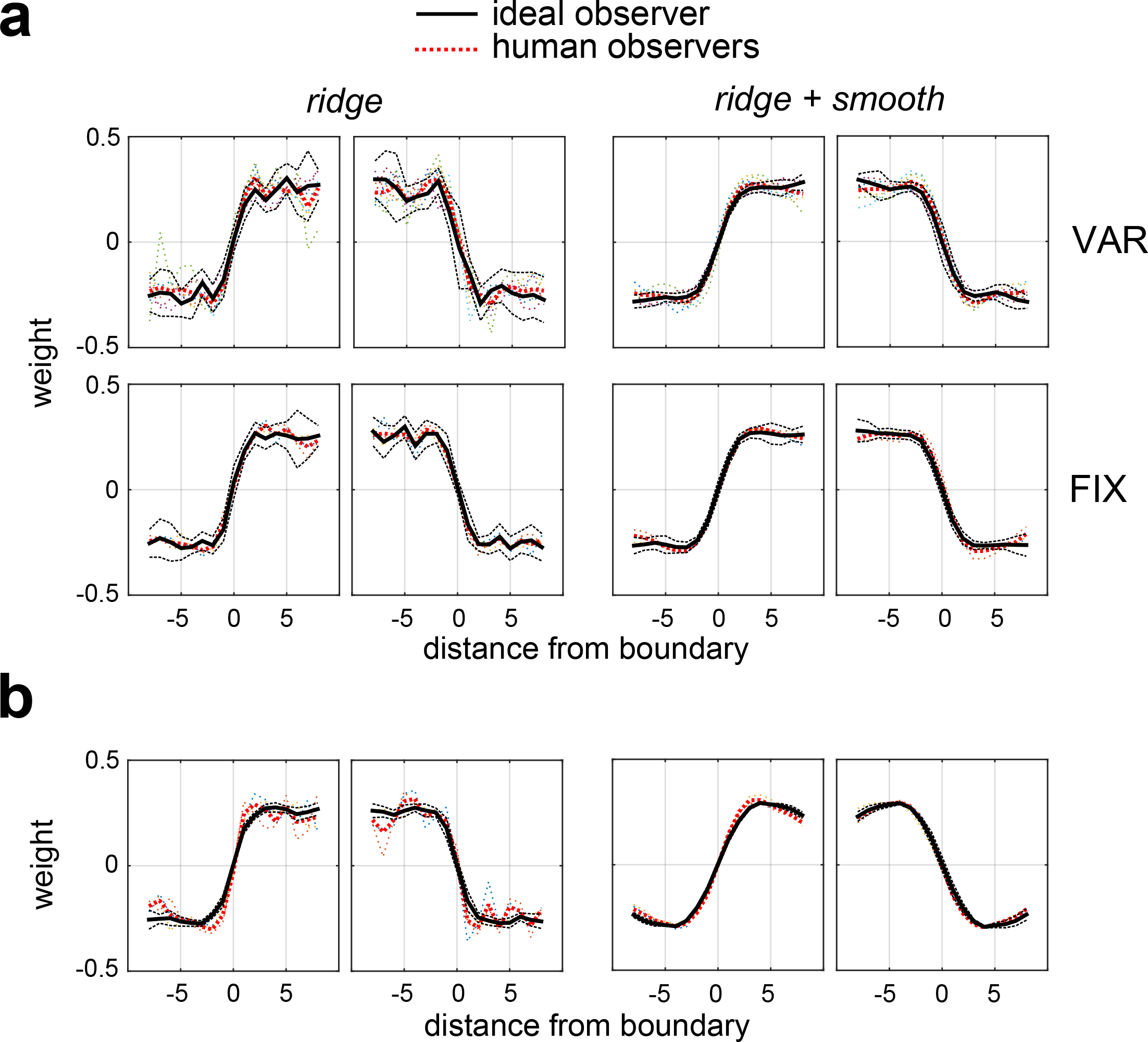
Normalized 1-D profiles of the average second-stage filter weights (30 bootstrapped re-samples). Thick black lines are for the ideal observer, thin black dashed lines indicate confidence intervals (+/- 1.96 SEM). Colored lines indicate normalized profiles for individual human observers. All models used AVG pooling. **(a)** Experiment 1-VAR (top) and Experiment 1-FIX (bottom). **(b)** Experiment 1-HFC (higher density of smaller micropatterns).

#### Second-stage filter nonlinearity

In Experiment 1, the best-fit expansive nonlinearity exponent α for the second-stage filters is greater than unity for all observers, for both the *ridge* and *ridge* + *smooth* priors. For Experiment 1-VAR with AVG sampling, median α values were around 2 (*ridge*: median = 2.15, min = 1.70, max = 2.85; *ridge* + *smooth*: median = 1.95, min = 1.70, max = 3.35, N = 7), and for Experiment 1-FIX we obtained median values closer to 1.5 (*ridge*: median = 1.50, min = 1.40, max = 1.60; *ridge* + *smooth*: median = 1.60, min = 1.60, max = 1.90, N = 3). Similar results were obtained for other downsampling methods (Supplementary Tables S2, S3), and the ideal observer was also best fit with an expansive nonlinearity (AVG; Experiment 1-VAR: *ridge*: 4.00; *ridge* + *smooth*: 3.30; Experiment 1-FIX: *ridge*: 2.20; *ridge* + *smooth*: 2.30). Plots of the second-stage nonlinearities for observers AMA, JJF are shown in Supplementary Fig. S11. This finding is consistent with previous studies suggesting an accelerating nonlinearity following the second-stage filters (see Discussion).

### Analysis of model accuracy

To better assess the meaningfulness of the above model-fitting results, it is important to make quantitative measures of the fitted models’ accuracy in predicting observers’ responses to novel stimuli. Such measures can also address the relative merit of different variants of the estimation procedure (*ridge* vs *ridge+smooth* prior) and of the model details, such as downsampling method (AVG, MAX, SUB) and classification rule (DET, STO). In this section we describe some of our analyses of model performance, and refer to other assessments in the Supplementary Material.

#### Psychometric function agreement

To the extent that the model architecture is appropriate and the data are sufficient to obtain accurate estimates of parameters, then the fitted model should be able to predict each observer’s responses. Consequently, we constructed psychometric functions of proportion correct on the orientation identification task as a function of the modulation depth across the boundary for individuals observers in Experiment 1-VAR - three representatives are shown in Fig. 6 (AVG downsampling). The model predictions (green, red) on the test datasets (which were not used to train models or optimize hyper-parameters) generally lie within or near the 95% binomial proportion confidence intervals (+/- 1.96 SEM) of observer performance (Fig. 6, blue dotted lines). However, the deterministic model (DET, left column) tends to over-predict performance at larger contrast modulations for some observers. Implementing the model stochastically (STO, right column) improved the model-observer agreement.

**Figure 6:**
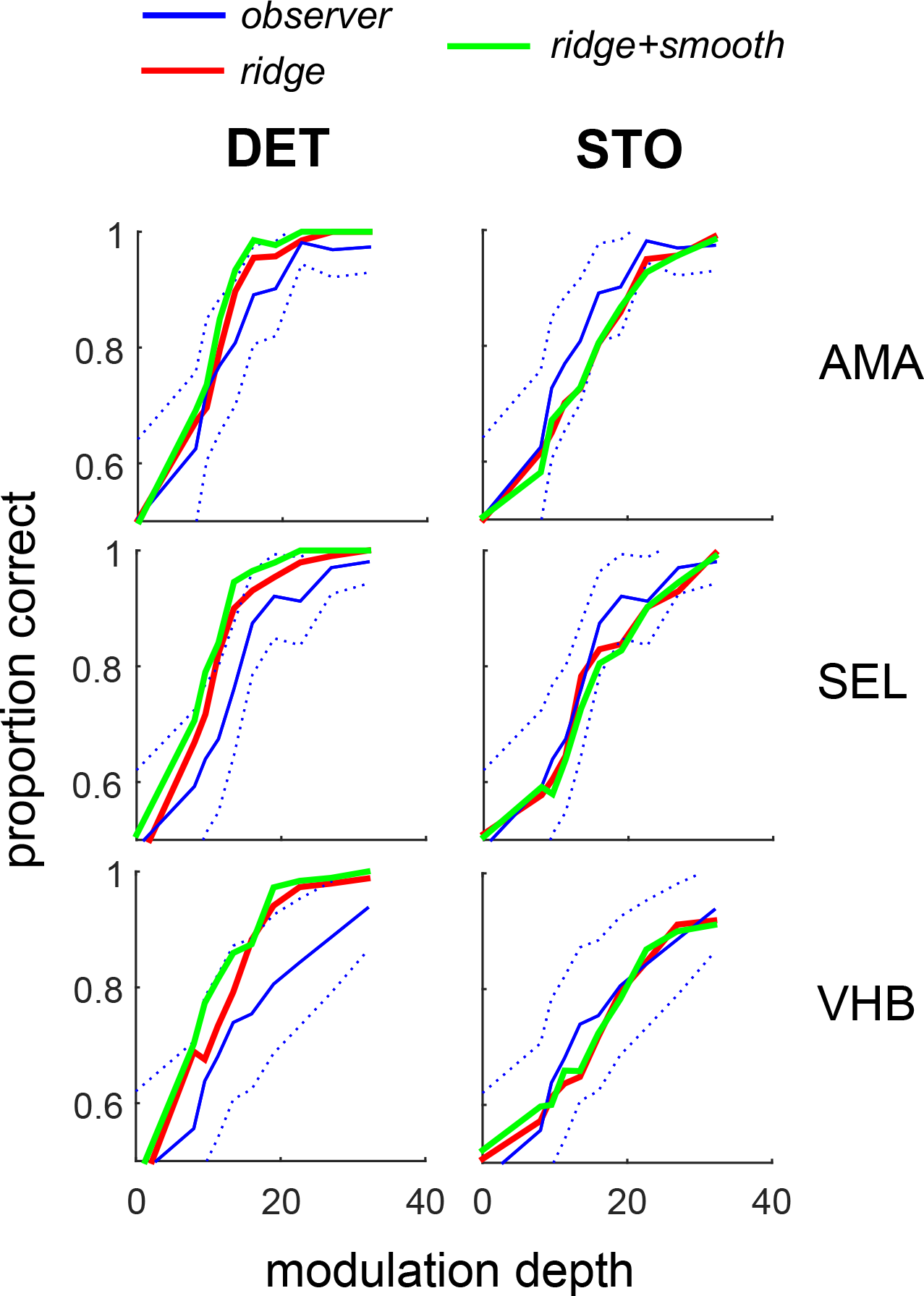
Psychometric functions for two versions of model (DET, STO), for three representative human observers (AMA, SEL, VHB) in Experiment 1-VAR. Blue lines denote human observer performance on the test set (stimuli not used for model estimation) as a function of modulation depth, together with 95% binomial proportion confidence intervals (+/- 1.96 SEM, blue dashed lines). Red and green lines denote model predictions for test stimuli for different Bayesian priors (red: *ridge*; green: *ridge* + *smooth*). Left column shows model operating deterministically (DET), right column shows model operating stochastically (STO).

#### Overall performance agreement

Another way to assess how well fitted models account for observer responses is to compare their overall performance (i.e. proportion correct averaged over all modulation depths tested) with observer performance. Such an analysis succinctly characterizes model accuracy for each observer and model variant with a single number (as opposed to an entire psychometric function). Fig. 7 shows observer and model performance in Experiment 1-VAR for each individual test fold on every observer (7 observers, 4 test folds each = 28 total folds). The deterministic model with AVG pooling (AVG-DET, Fig. 7a, top) somewhat tended to over-predict observer performance, with the majority (*ridge*: 18/28; *ridge* + *smooth*: 23/28) of test folds having significantly different (binomial proportion difference test, 95% CI) proportions correct (Fig. 7b, top). Despite the systematic over-prediction of observer performance (by about 6-8%), we still observe significant correlations between predictions and actual performance across observers (*ridge*: *r* = 0.983, *p* < 0.001; *ridge* + *smooth*: *r* = 0.972*, p* < 0.001, N = 7) and folds (*ridge*: *r* = 0.872, *p* < 0.001; *ridge* + *smooth*: *r* = 0.806*, p* < 0.001, N = 28).

**Figure 7:**
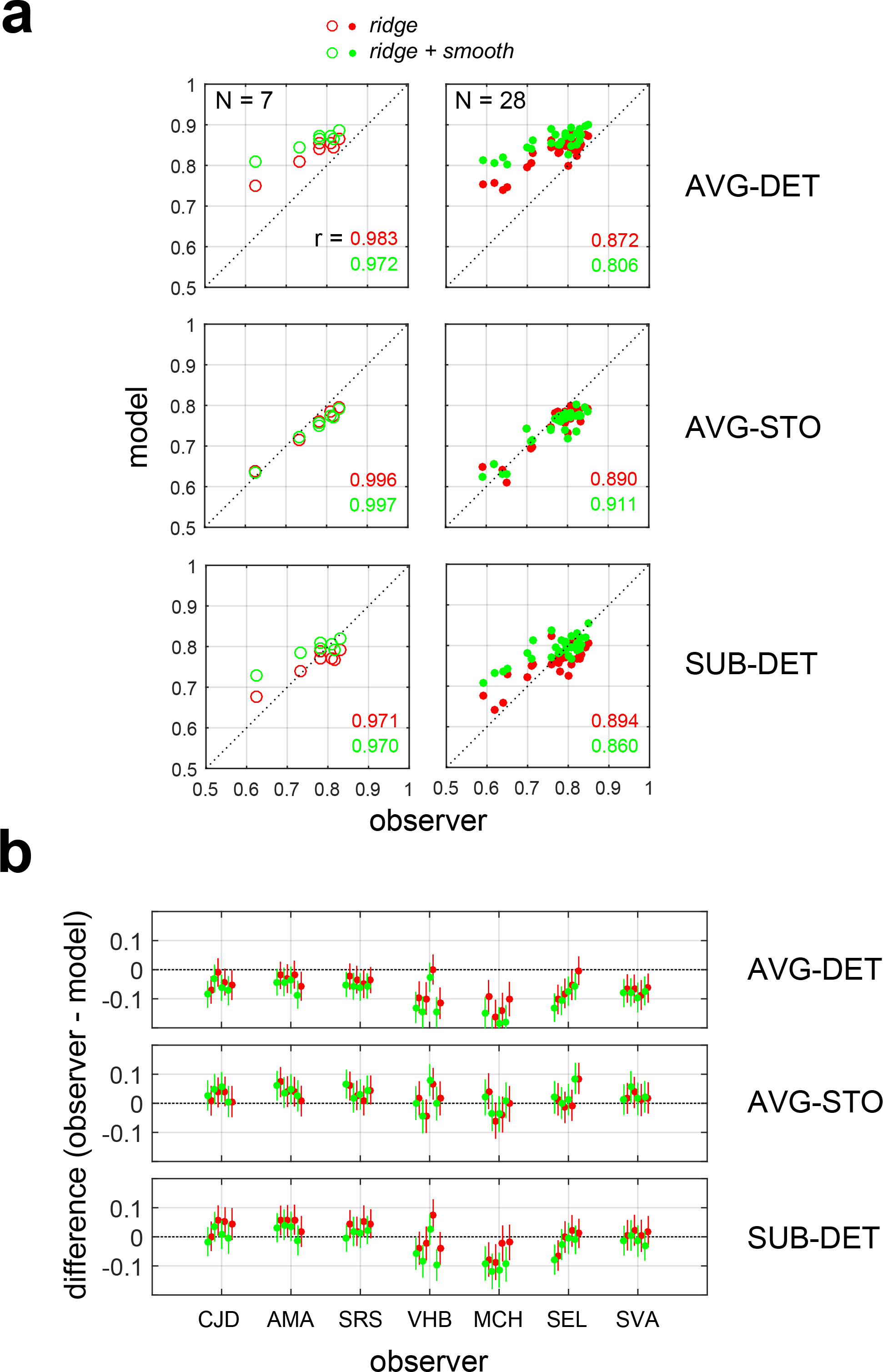
Analysis of model accuracy for Experiment 1-VAR. (**a**) Plots of model vs. observer performance (proportion correct), averaged across observers (left sub-panels, circles, N = 7) or test folds (right sub-panels, solid symbols, N = 28). Observer and model performance were compared on a set of novel test stimuli (N = 500) not used for model estimation or hyper-parameter optimization. *Top:* Deterministic model with AVG (average) downsampling (AVG-DET). Correlation coefficients (r values) are color coded for each choice of Bayesian prior (red: *ridge*; green: *ridge* + *smooth*). *Middle:* Stochastic model with AVG downsampling (AVG-STO). *Bottom:* Deterministic model with SUB downsampling implemented with subsampling (SUB-DET). (**b**) Difference between observer and model performance for each individual test fold (4 folds per observer) for all models shown in (a). Lines show 95% confidence intervals of the difference (binomial proportion difference test). Colors indicate different choices of Bayesian prior (red: *ridge*; green: *ridge* + *smooth*).

Implementing the classification stochastically produced much better agreement (AVG-STO: Fig. 7a, middle), with relatively few (*ridge*: 4/28; *ridge* + *smooth*: 6/28) test folds having significantly different proportions correct (Fig. 7b, middle). As before, strong correlations between model and observer performance are seen across observers (*ridge*: *r* = 0.996, *p* < 0.001; *ridge* + *smooth*: *r* = 0.997, *p* < 0.001, N = 7) and folds (*ridge*: *r* = 0.890, *p* < 0.001; *ridge* + *smooth*: *r* = 0.911, *p* < 0.001, N = 28). For the deterministic (DET) model, performance was more accurate for subsampling (SUB-DET: Fig. 7a, bottom) than averaging downsampling (AVG-DET: Fig. 7a, top) or MAX-DET downsampling (not shown). In this case, many fewer folds (*ridge*: 10/28; *ridge* + *smooth*: 8/28) had significantly different proportions of correct responses (Fig. 7b, bottom). Similar results were obtained for Experiment 1-FIX (Supplementary Fig. S13).

The over-prediction of observer performance for the DET model with AVG + MAX pooling is due to the DET mode of operation not accounting for internal noise on a trial-by-trial basis. By contrast, this noise is accounted for when these model is operating in STO mode, since the model is explicitly optimized to predict the proportion of correct responses at each level. The degradation in model performance for SUB-DET pooling (leading to better agreement with the observer) may be due to the fact that this model is getting less information from the first-stage filters than AVG or MAX pooling, which integrate over all first-stage filters instead of a relatively small subset of them. In an additional analysis of Experiment 1-FIX, we systematically varied the downsampling rate for all three pooling rules (Supplementary Fig. S12). We find that when operating in DET mode, for low rates (8×8) the SUB model drastically under-predicts observer performance, while for higher rates (16×16, 22×22) the SUB model performance improves and more closely matches observer performance. By contrast, the we see in Supplementary Fig. S12 that performance of the DET mode AVG and MAX models consistently over-predicts performance for all down-sampling rates, since regardless of down-sampling rates both pooling rules integrate over all of the first-stage filters rather than a limited subset, making them highly robust to random variations in micropattern placement.

#### Double-pass analysis

To obtain an upper bound on the performance of the model, we carried out a “double-pass” analysis (Neri & Levi, 2006; Neri, 2010; Hasan, Joosten, & Neri, 2012) by testing observers twice on the same set of stimuli, and comparing model-observer agreement (proportion of trials with consistent responses) on the double-pass set to observer-observer agreement between the two passes. For 5 observers in Experiment 1-VAR, a fold of 1000 stimuli was used as the double-pass set, and the model was trained on the remaining 3000 stimuli. The results (Fig. 8) show that the AVG-DET or MAX-DET models agree with the observer on the two passes as well as the observer is self-consistent on the two passes. For AVG-DET, both priors give a small, statistically insignificant (rank-sum test) median difference (*ridge*: median difference = 0.022, *p* = 0.063; *ridge* + *smooth*: median difference = 0.009, *p* = 0.313, N = 5), with most or all of the (binomial proportion difference) confidence intervals bracketing zero (*ridge*: 4/5; *ridge* + *smooth*: 5/5). Similarly, MAX-DET gives small median differences (*ridge*: median difference = 0.018, *p* = 0.125; *ridge* + *smooth*: median difference = 0.02, *p* = 0.313, N = 5) with most confidence intervals (*ridge*: 4/5; *ridge* + *smooth*: 4/5) including zero. However, SUB-DET (Fig. 8a, bottom panels) gives somewhat larger differences (*ridge*: median difference = 0.052, *p* = 0.063; *ridge* + *smooth*: median difference = 0.024, *p* = 0.063, N = 5), although they fail to reach statistical significance. In the SUB-DET case with the *ridge* prior (red), 0/5 confidence intervals include zero, while 3/5 do so for *ridge* + *smooth* (green). Implementing the decision rule stochastically (AVG-STO) yielded somewhat poorer double-pass agreement (Fig. 8b), although the difference fails to reach significance across the set of observers (*ridge*: median difference = 0.075, *p* = 0.063; *ridge* + *smooth*: median difference = 0.088, *p* = 0.063, N =5). For AVG-STO, we find that for both priors, 0/5 confidence intervals contain zero.

**Figure 8:**
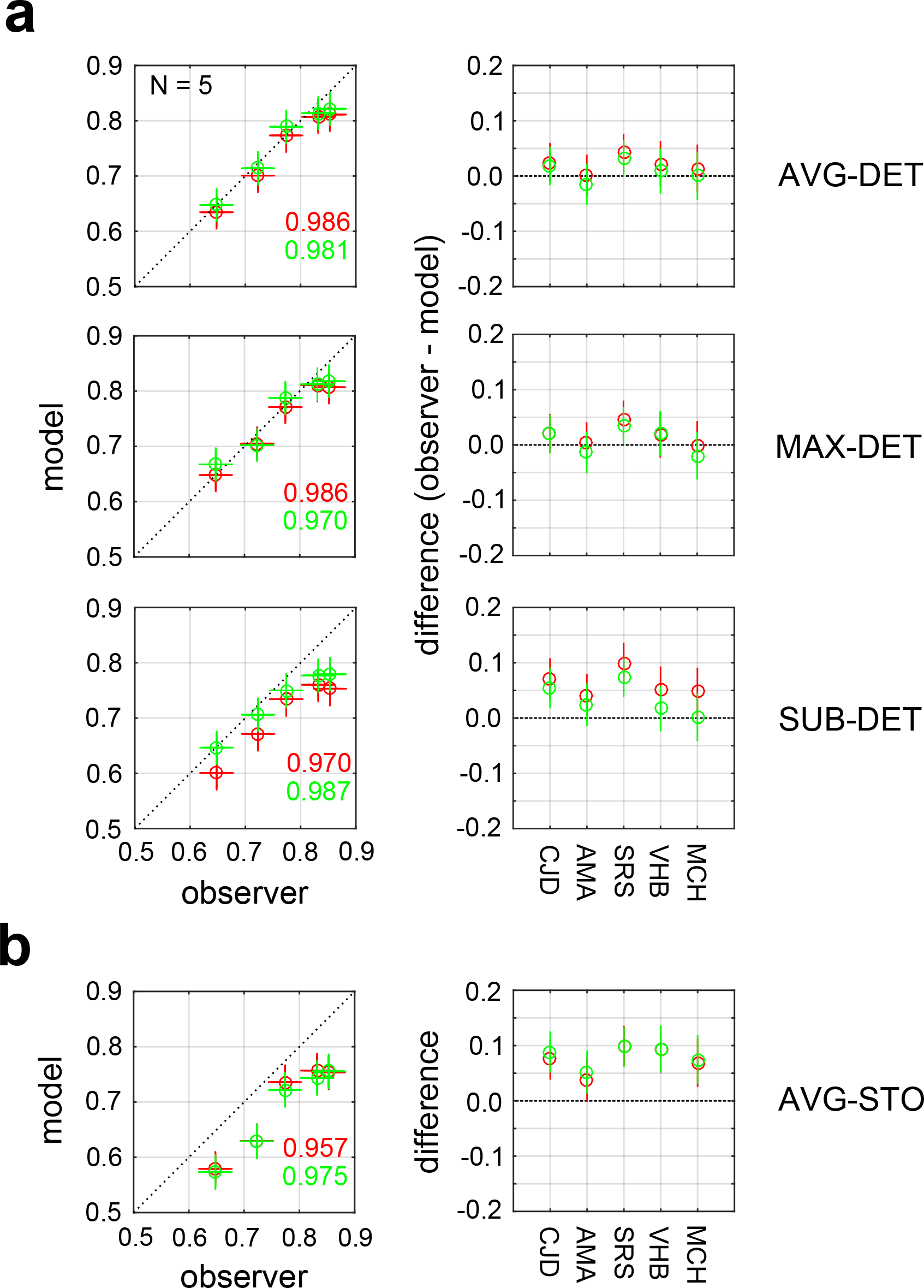
Double-pass analysis for 5 observers in Experiment 1-VAR. (**a**) Models operating in deterministic (DET) mode for three different downsampling rules. *Left panels:* Observer-observer (horizontal axis) and observer-model (vertical axis) proportion agreement on the double-pass set. Error bars denote 95% binomial proportion confidence intervals. *Right panels*: Difference between observer-observer and observer-model proportion agreement for each of 5 observers. Error bars denote 95% binomial proportion difference confidence intervals. (**b**) Same as (a), but for model with AVG downsampling operating in stochastic mode (AVG-STO).

Why does the stochastic (STO) decision rule (with AVG/MAX pooling) lead to better prediction of observer performance (Fig. 7), and yet worse double-pass agreement (Fig. 8) than the deterministic (DET) rule? To understand why, suppose for a given right-oblique stimulus the observer gave the response (R) on the first trial, and due to internal noise is 80% likely to respond R on the second trial, yielding a self-consistent response (R, R) on both passes. Suppose that for this stimulus the output of the model is P(R) >0.5. Then the DET model will always return R, and therefore be in agreement with the observer’s choice on the second pass 80% of the time. In contrast, the STO model will return R on only a subset of the trials, which will be uncorrelated with the observer’s responses. Therefore, the STO model will have a lower rate of double-pass agreement, even if it better matches the overall proportion correct.

#### Decision variable correlation analysis

Although a strong model-observer agreement for the percentage correct is consistent with accurate modeling of the observer’s visual processing, it is not fully compelling. The model and observer might both choose category A or B the same proportion of trials, without necessarily exhibiting any correlation between observer and model responses on a trial-by-trial basis. This situation could arise, for example, if the observer and model were utilizing different stimulus features for their decisions. To more thoroughly quantify model performance, we implemented a *decision variable correlation* (DVC) analysis, which estimates the trial-by-trial correlation between a model’s decision variable and the observer’s hypothesized decision variable (Sebastian and Geisler, 2018). This method extends standard signal detection theory by hypothesizing that the joint distribution of model and observer decision variables for each stimulus category (A = left-oblique/L, B = right-oblique/R) can be modeled as bivariate Gaussian distributions having correlation parameters ρ_A_, ρ_B_. Using the proportions of trials on which the observer and model decisions agree for each stimulus category, ρ_A_ and ρ_B_ are estimated using a simple maximum likelihood procedure (Sebastian & Geisler, 2018).

DVC analysis is most accurate (as measured by bootstrapped estimates of SEM) for large numbers of trials at a constant stimulus level sufficiently near threshold to give consistent proportions of the four logical possibilities (model says A/B, observer says a/b) (Sebastian & Geisler, 2018). Therefore we applied DVC analysis to Experiment 1-FIX, which entailed thousands of trials (6000) at a single stimulus level for each of three observers (CJD, MAK, JJF). The model was trained on 5000 trials and the DVC computed using the remaining 1000 trials, for 6 choices of hold-out fold. Since the two boundary orientations provide independent estimates of the DVC, this provided 12 estimates of DVC from each observer. Fig. 9 shows the resulting DVC values (plotted in rank order within each set, for graphical clarity) measured for AVG downsampling and both choices of prior. These estimates of DVC are overwhelmingly positive and significantly larger than zero.

**Figure 9:**
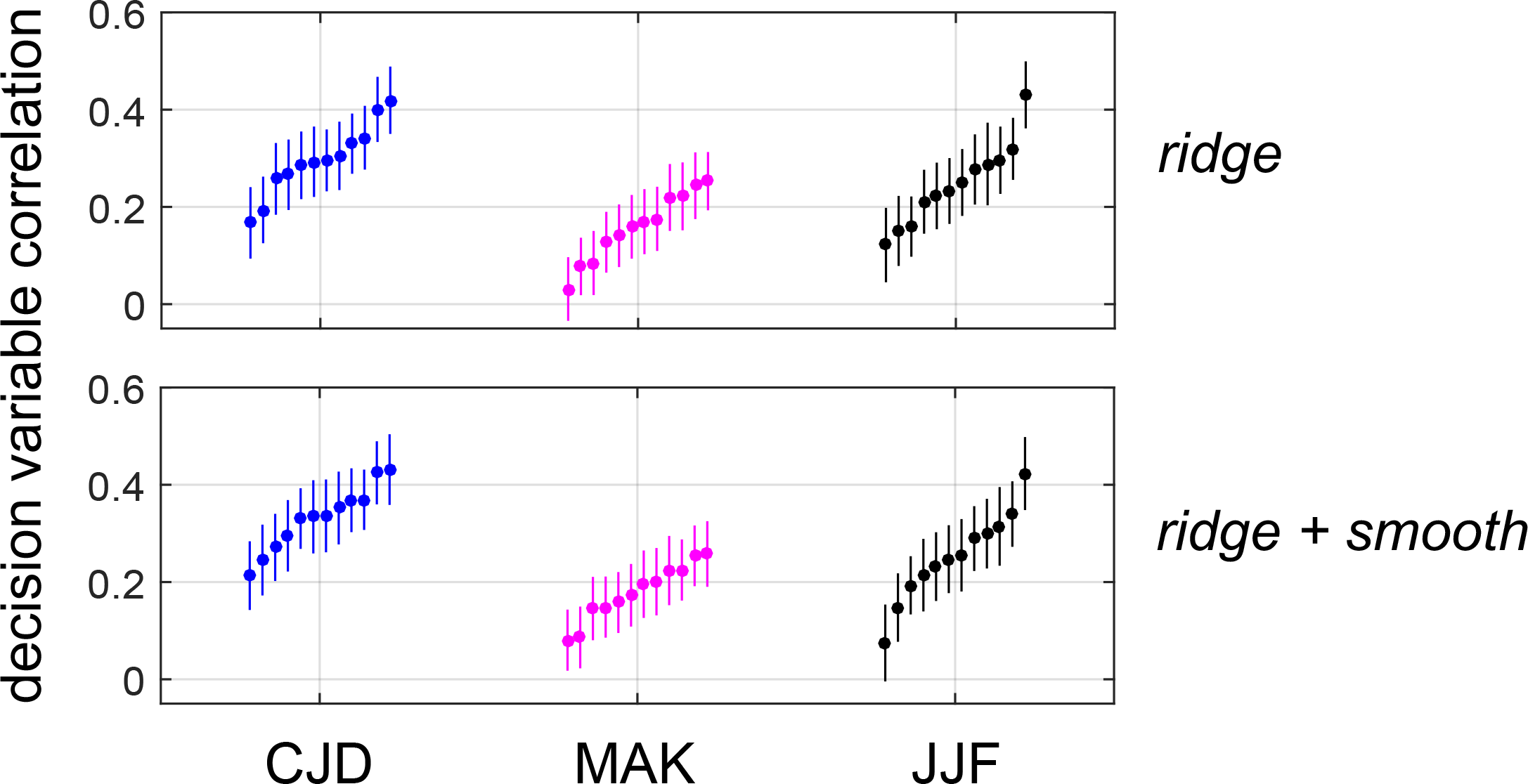
Decision variable correlation analysis. Decision variable correlation (DVC) computed on 6 test folds (two measurements per fold) for three observers (CJD = blue, MAK = magenta, JJF = black) in Experiment 1-FIX for both priors. DVC values for each observer are sorted by magnitude for visual clarity. Thin lines denote +/-1 SEM (200 bootstrap re-samples).

For AVG downsampling, 32/36 (*ridge*) and 33/36 (*ridge* + *smooth*) estimates of DVC are significantly (> 1.96 SEM) larger than zero, with DVC mean +/- SD of 0.234 +/- 0.094 (*ridge*) and 0.254 +/- 0.095 for (*ridge* + *smooth*). MAX downsampling gives similar results, where 33/36 (*ridge*) and 35/36 (*ridge* + *smooth*) estimates of DVC are significantly greater than zero, with DVC mean +/- SD of 0.245 +/- 0.094 (*ridge*) and 0.263 +/- 0.086 (*ridge* + *smooth*). Interestingly, SUB downsampling gives weaker correlations with only 20/36 (*ridge*) and 24/36 (*ridge* + *smooth*) estimates of DVC significantly greater than zero, and DVCs have mean +/- SD of 0.147 +/- 0.082 (*ridge*) and 0.164 +/- 0.085 (*ridge* + *smooth*).

Therefore, we see that not only are the models and observers in agreement about overall proportions of each category, but they are also correlated in their trial-by-trial decisions, consistent with the interpretation that the models and human observers are performing similar kinds of processing.

#### Ideal-observer analysis

To compare the spatial summation and task performance of human observers to those of an ideal observer, we trained the model on the true stimulus category labels rather than observer classifications of these stimuli. Notably, the resulting ideal observer models make use of a region-based spatial summation similar to that of human observers (Fig. 3c). Examining this more systematically, a comparison of human performance to that of the deterministic (16×16 AVG-DET) ideal observer in Experiment 1-VAR (Supplementary Fig. S14) revealed that across 6 observers analyzed, most confidence intervals (binomial proportion difference test, 95% confidence interval) did not contain zero (*ridge*: 6/24; *ridge + smooth:* 3/24), with those containing zero coming mainly from two observers (AMA, SRS). In contrast, for the stochastic ideal observer (AVG-STO) many more confidence intervals contained zero (*ridge*: 11/24; *ridge* + *smooth*: 10/24), suggesting a better agreement with observer performance. Similar results were obtained for the 3 observers tested in Experiment 1-FIX (Supplementary Fig. S14). Comparing human to deterministic ideal observer (AVG-DET) performance, most confidence intervals did not contain zero (*ridge*: 5/18; *ridge* + *smooth*: 3/18), while for the stochastic ideal observer most did contain zero (*ridge:* 12/18; *ridge* + *smooth*: 15/18). On the whole, this result is similar to our findings for the models fitted to observer data – i.e. a general tendency for the AVG or MAX models operating deterministically to over-predict performance, while these models operating stochastically seemed to agree reasonably well.

#### Correlations between observer performance and model parameters

The different human observers exhibited substantial individual differences in overall performance as measured by proportion correct (Fig. 7), raising the question whether there might be corresponding variations in the fitted model parameters. We assessed this possibility for the results in Experiment 1-VAR, using the 16×16 AVG model.

Although the nonlinearity was always expansive (exponent α > 1.0), there was no compelling evidence across the set of individuals that the magnitude of α explained individual variations in performance, based on Pearson’s *r* (*ridge*: *r* = -0.458, *p* = 0.302; *ridge + smooth: r* = -0.787, *p* = 0.036, N = 7). The significant result for the *ridge* + *smooth* case was due to the presence of an outlier - computing these correlations using Spearman’s rho revealed no significant effect (*ridge: rho* = -0.357, *p* = 0.444; *ridge* + *smooth*: *rho* = -0.214, *p* = 0.662). Results were similar for MAX and SUB models (Supplementary Table S5).

However, individual variability in observer performance was very strongly related to the magnitudes of the fitted second-stage filter weights. That is, correlating the sum of the norms of the filter weights with overall performance across the 7 observers yielded strong, significant positive correlations (*ridge*: *r* = 0.970, *p* < 0.001; *ridge* + *smooth*: *r* = 0.993, *p* < 0.001), with similar results for MAX, SUB models (Supplementary Table S6). Consistent with this, the ideal observer had a larger filter norm (*ridge*: 5.67; *ridge* + *smooth*: 5.37) than the human observers (*ridge*: median = 4.30, min = 2.36, max = 4.59; *ridge + smooth:* median = 4.04, min = 2.27, max = 4.34, N = 7), as revealed by a sign-rank test (*ridge: p* = 0.018; *ridge + smooth: p* = 0.018).

In our model, the inverse internal noise level is represented implicitly by the magnitudes (vector norms) of the second-stage filters (see Methods for details). Therefore, we should expect observers with larger second-stage filter norms to also have better double-pass agreement, which is a function of internal noise. We find that there are indeed strong, significant positive correlations between filter norm and double-pass agreement for all pooling rules and choices of prior (AVG - *ridge*: *r* = 0.954, *p* = 0.012, *ridge* + *smooth*: *r* = 0.942, *p* = 0.017; MAX - *ridge*: *r* = 0.964, *p* = 0.008, *ridge* + *smooth*: *r* = 0.956, *p* = 0.011; SUB - *ridge*: *r* = 0.940, *p* = 0.017, *ridge* + *smooth*: *r* = 0.943, *p* = 0.016). Taken together with significant positive correlations between observer performance and double-pass agreement (*r* = 0.949, *p* = 0.014), we conclude that variations in filter norm magnitudes across observers are related to variations in internal noise levels across observers, and this variability in internal noise levels contributes to difference in observer performance.

#### Summary and Interpretation

From our analysis of model accuracy, we may draw several broad conclusions. Firstly, accuracy is not greatly affected by choice of prior, with similar results obtained using the *ridge* and *ridge* + *smooth* priors. Also, from our analyses we cannot favor a particular choice of pooling rule among those tested (AVG, MAX, SUB). Almost identical overall results are obtained using AVG and MAX pooling, and although in some cases we see lower accuracy using SUB pooling, our results suggest this can be ameliorated by simply increasing the down-sampling rate (Supplementary Fig S12). In Experiment 1-VAR, where the stimulus levels were fixed for all observers (rather than set at their individual thresholds), we observed great variability between observers in both task performance and double-pass agreement. This variability is reflective of differences in internal noise across observers, and is captured by our model.

### Experiment 2: Boundary orientation discrimination

The first experiment demonstrated that human observers employ region-based processing for the orientation identification task. To see if our modeling method could also reveal situations where observers employ edge-based processing, we utilized a fine orientation discrimination task (Fig. 1c, right). The orientation of a second-order boundary was perturbed slightly (JND at 32% contrast modulation – see Methods) from vertical, and observers judged the direction of perturbation (i.e. slightly left- vs right-oblique). In one variant of this task (Experiment 2-VAR), contrast modulation was varied in 9 logarithmic steps from 16% to 64% centered at 32% (plus zero modulation) to vary the task difficulty. In a second variant (Experiment 2-FIX), contrast modulation was fixed at 32%. In both variants of Experiment 2, the perturbation of orientation was fixed near its JND for each observer at 32% contrast modulation (CJD: 7 deg., JJF: 6 deg., VHB: 6 deg.). In the data analysis, a higher downsampling factor (22 × 22) was used to better reveal fine spatial details, and only AVG pooling was considered, since Experiment 1 demonstrated substantial robustness of spatial summation to pooling rules (Supplementary Figs. S8, S9).

Fig. 10 shows the second-stage filter weights from an observer (JJF) tested in both Experiment 2-VAR (Fig. 10a) and Experiment 2-FIX (Fig. 10b). Unlike in Experiment 1, here observers employ an “edge-based” summation, assigning substantially more weight to texture regions near the boundary. Bootstrapped significance plots for individual weights are shown in Supplementary Fig. S15, and consistent results from other observers are shown in Supplementary Fig. S16. The edge-based processing employed by these observers is very similar to that obtained from an ideal observer (Supplementary Fig. S17). As with Experiment 1, we find a reasonably strong agreement between model predictions and observed performance on sets of novel data not used for model parameter estimation (Supplementary Fig. S18), and find an expansive second-stage filter nonlinearity for Experiment 2 (Supplementary Table S7). Taken together, Experiments 1 and 2 show that our modeling methodology can reveal different kinds of spatial processing that observers may employ for different kinds of psychophysical tasks.

**Figure 10:**
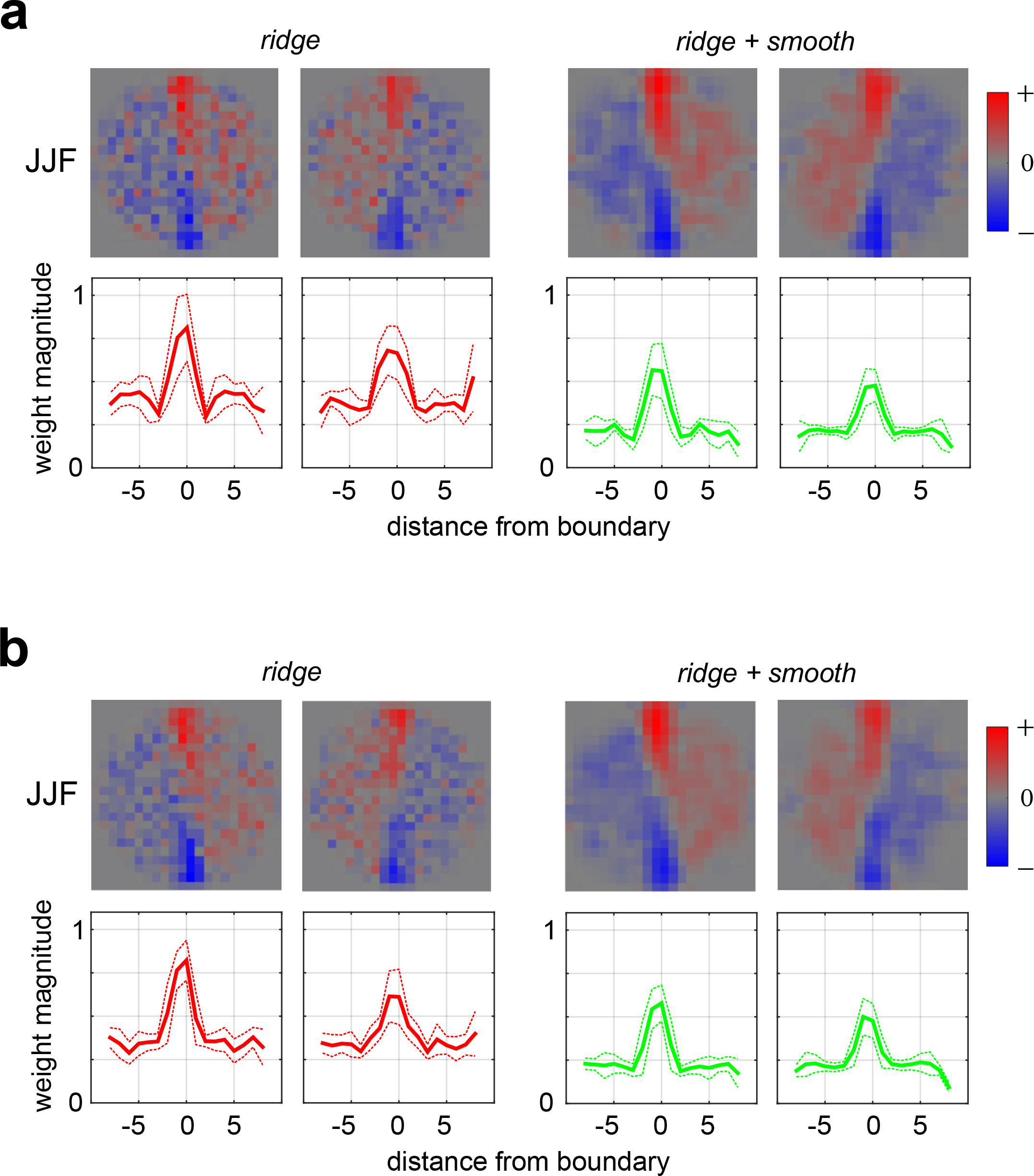
Second-stage filter weights for one representative observer, JJF, from Experiment 2 (orientation discrimination, Fig 1c, right). **(a)** Experiment 2-VAR. *Top:* Estimated 2-D weight maps averaged over 4 training folds. *Bottom:* 1-D profiles showing mean weight magnitude (thick lines), averaged over 30 bootstrapped re-samplings. Thin dashed lines show +/- 1 SEM. **(b)** Same as (a), but for Experiment 2-FIX.

### Experiment 3: Integrating contrast modulation across orientation channels

In Experiments 1 and 2, we considered a particular FRF model architecture and focused on characterizing the spatial processing that observers employ in different second-order vision tasks. However, one of the most powerful applications of this methodology is model comparison - i.e. comparing the ability of alternative models, embodying different assumptions about an FRF model architecture, to account for psychophysical data. Indeed, a major focus of much of the second-order vision literature has been to design experiments whose express goal is to test competing hypotheses of FRF architecture (Motoyoshi & Kingdom, 2003; Motoyoshi & Nishida, 2004; Kingdom, Prins & Hayes, 2003; Motoyoshi & Kingdom, 2007; Westrick & Landy 2013).

We applied our method to ask how second-stage filters for contrast modulation integrate orientation modulations across multiple kinds of first-stage filters or channels. In many previous computational studies of second-order processing, the FRF models were comprised of second-stage filters, each of which only analyzes a single orientation/spatial frequency channel (Wilson 1993; Landy & Oruc, 2002; Zavtiz & Baker 2013), with subsequent “late summation” of their signals. However, other psychophysical studies have suggested “early summation” FRF models, in which second-stage filters integrate contrast modulation across texture channels (Motoyoshi & Kingdom, 2003; Motoyoshi & Nishida, 2004; Kingdom, Prins, & Hayes, 2003).

To address this question systematically, we used a boundary orientation identification task as in Experiment 1, but with the carrier pattern containing two orthogonal orientations of micropatterns (Fig. 1a, bottom), which would be processed by distinct first-stage orientation-selective channels. Since the nature of spatial summation is not critical to this question, we used a coarser downsampling factor (12 × 12), and as above, only AVG pooling was used. We considered two models of how information is integrated by the second-stage filters. In a “late summation” Model 1 (Fig. 11a, top), each first-stage channel is analyzed by its own pair of second-stage filters (L/R oblique). Alternatively, in an “early summation” Model 2 (Fig. 11a, bottom), each second-stage filter receives inputs from both first-order channels. As in Experiments 1 and 2, we find that both the models are in reasonably good agreement with observer performance on the task (Supplementary Fig. S20). Also there was reassuring consistency in that both the fitted models exhibited similar region-based weight maps, with an expansive second-order nonlinearity (Supplementary Table S8).

**Figure 11:**
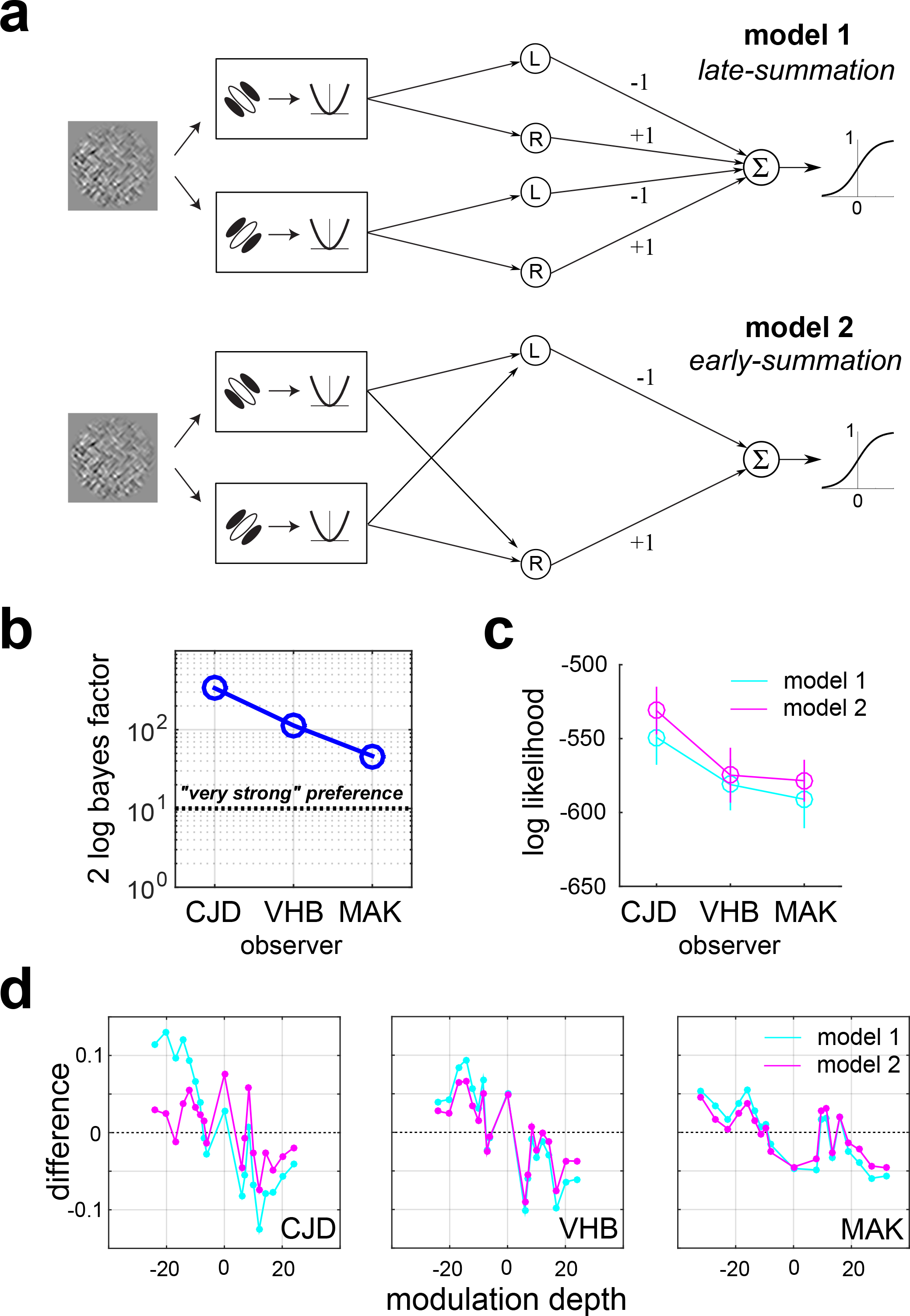
Model comparison for Experiment 3, boundary orientation identification with textures composed of two kinds of micropatterns. **(a)** Two competing models of FRF architecture fit to psychophysical trial data. *Top:* Model 1 (“late summation”) assumes that each first-stage orientation channel is analyzed by its own pair of (L/R) second-stage filters and then pooled. *Bottom:* Model 2 (“early summation”) assumes that each (L/R) second-stage filter integrates over both first-stage channels. (**b**) Bayesian model comparison making use of all data reveals a strong consistent preference (as measured by the Bayes factor – see text) for Model 2 (“early summation”) for all three observers (blue line and symbols). Thick black line indicates the conventional criterion for a “very strong” preference for Model 2. (**c**) Bootstrapping analysis, in which model likelihood is evaluated on novel data not used for model training, also reveals a preference for Model 2. Plotted points indicate mean +/- SEM for 50 bootstrapped samples. (**d**) Difference between predicted and observed proportions where observer chooses “right oblique” for all observers and both models. Negative modulation depths indicate left-oblique stimuli.

When comparing Models 1 and 2 using standard model comparison techniques such as the Bayes Information Criterion, or BIC (Schwarz, 1978; Bishop 2006), there is no need to correct for overfitting since both models have the same number of parameters. This is because when parameter space dimensionality is equal, the complexity-penalizing terms of the BIC cancel, so that Δ_*BIC*_= ln *p*_2_(*D*) − ln *p*_1_(*D*), where *p*_*i*_(*D*) denotes the likelihood of the data (*D*) under model *i* evaluated at the MAP estimate of the model parameters. That is, when two models are of equal complexity and equally likely *a priori*, we simply choose the one with higher log-likelihood of the data. The difference in log-likelihoods (Δ_*BIC*_) is identical to the natural log of the Bayes factor 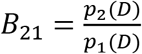 . Standard effect size conventions state that when 2 ln *B*_21_ > 10 (corresponding to > 150-fold higher probability of Model 2), the preference is considered to be “very strong” (Kass & Raferty, 1995). We see in Fig. 11b that for all three observers tested in Experiment 3, the evidence (measured by 2 ln *B*_21_) in favor of the orientation opponent model (Model 2) exceeds the standard criterion (10) for a very strong preference. Ideal observer analysis also revealed a very strong preference for the orientation-opponent model fit to stimulus category labels (2 ln *B*_21_ = 478.4). With a large number of trials, a strong preference for one binomial response model does not require a huge difference in predicting observer performance (Appendix A). We see in Fig. 11d that although Model 2 very consistently does a better job of matching observer performance than Model 1, that the differences between predicted proportions correct are fairly small.

We also wanted to compare the predictive abilities of the two models using a cross-validation approach, by evaluating their agreement with observer performance on data not used for training. Therefore we performed a bootstrap analysis in which both models were fit on randomly selected samples of 4000 stimuli and tested on the remaining 1000 (50 bootstraps). To optimize the generalization performance, each model was trained using the value of the ridge regression hyperparameter λ which yielded the best generalization (Supplementary Figs. S19, S21). The results in Fig. 11c show a consistent preference across observers for Model 2 (magenta line). Performing a 3 × 2 factorial ANOVA (3 observers, 2 models, 50 observations each) on the test set log-likelihoods revealed significant main effects of both observer (*F*_2,294_ = 201.085, *p* < 0.001) and model (*F*_2,294_ = 39.811, *p* < 0.001), but no significant interaction (*F*_2,294_ = 3.001, *p* = 0.051). In summary, both analysis approaches (Figs. 11b, c) show a strong and consistent preference for the early summation version (Model 2).

## Discussion

Second-order stimuli present a challenge for computational models of visual processing, as they do not contain spatial variations in stimulus intensity and thus cannot be detected by simple linear filtering operations (Baker, 1999). Consequently, standard “classification image” techniques (Ahumada, 1996; Murray, 2011), which assume a linear spatial summation model, are not readily applicable to studies of second-order visual processing. Here we define a simple hierarchical neural network implementing a Filter-Rectify-Filter (FRF) model of second-order visual processing (Bovik, Clark, & Geisler, 1990; Malik & Perona, 1990; Bergen & Landy, 1991; Sutter, Beck, & Graham, 1989; Landy & Graham, 2004). We estimate its parameters directly from psychophysical trial data, and show that it can accurately predict observer performance on novel test data not used for training. We demonstrate the utility of this modeling approach by testing whether observers segment contrast-defined boundaries using an edge- or region-based processing, finding support for region-based segmentation in a boundary orientation-identification task (Experiment 1), and edge-based processing for a boundary discrimination task (Experiment 2). In addition, we show how this methodology can adjudicate between alternative FRF model architectures (Experiment 3).

### Model complexity vs. biological realism

Psychophysical system identification is critically limited by the modest amount of data that can be collected in an experiment. Although constraints or priors can help alleviate the demand for data (Knobloch & Maloney, 2008a, b; Mineault et al., 2009), it is also often necessary to fit simplified models requiring fewer parameters, which omit many known biological details. Here we discuss several modeling simplifications which may need to be addressed when extending this methodology to more complicated second-order stimuli and tasks.

#### First-stage filters

In FRF models like those considered here, the initial stage of filtering and rectification is meant to model computations most commonly believed to be taking place in visual area V1 (Baker & Mareschal, 2001). We modeled the first-stage filters using a standard energy model of V1 complex cells with normalization (Adelson & Bergen, 1985; Heeger 1992). Although for our stimuli a front-end comprised entirely of oriented energy filters was perfectly adequate, for many texture segmentation tasks it may be necessary to model half-wave rectified filters (both even and odd phase) as well. For instance, human observers can segment second-order images using cues of carrier contrast phase and polarity (Malik & Perona, 1990; Hansen & Hess, 2006), which is not possible using only energy filters, whose responses are invariant to phase and polarity. Likewise, it may also be desirable to include center-surround “difference-of-Gaussian” (DOG) filters as well, since early visual cortex has many neurons with non-oriented receptive fields (Hubel & Livingstone, 1985; Ringach, Shapley, & Hawken, 2002; Talebi & Baker, 2012). Human psychophysics has indicated that non-oriented filters may potentially serve as a mechanism for polarity-based segmentation (Motoyoshi & Kingdom, 2007), and computer vision work on natural image segmentation found it advantageous to include non-oriented as well as oriented filters (Martin, Fowlkes and Malik, 2004).

In Experiments 1 and 2, we implemented only a single spatial frequency and orientation channel matched to the carrier, and in Experiment 3 we only considered two such channels. This simplification neglects the possible contributions of off-orientation and off-spatial frequency channels to task performance. Indeed, previous psychophysical studies have challenged the notion that second-order mechanisms are narrowly tuned for a single first-order channel, with many studies suggesting broad bandwidth for carrier spatial frequency (Graham, Sutter, & Venkatesan 1993; Prins & Kingdom, 2002; Westrick & Landy 2013). On the other hand, single unit neurophysiology has demonstrated neurons that are selective for the carrier spatial frequency of CM stimuli, both in macaque V2 (Li et al, 2014) and in cat Area 18 (Li & Baker, 2012).

#### Second-stage filters

The main focus of our study was demonstrating an approach to estimate the properties (shapes and nonlinearities) of a set of second-stage filters which detect higher-order visual features. For our task involving contrast-modulated boundaries (Fig 1c), the simplest possible model involves two second-stage filters representing the two potential stimulus categories, much like the model shown in Fig. 2. However, for psychophysical tasks involving more complex second-order stimuli such as natural occlusion boundaries (Fig. 1a, top) defined by multiple cues which differ on opposite sides of a boundary, it may be necessary to have both a wider variety of first-stage filters as well as multiple kinds of second-stage filters sensitive to different feature combinations (Fig. 12). In addition, it may be pertinent to allow direct connections between the first stage of filtering (albeit for much lower spatial frequencies) and the output (decision) stage, since natural region boundaries contain both first- and second-order cues (Schofield, 2000; Johnson & Baker, 2004; Martin et al., 2004; DiMattina et al., 2012) which can be integrated in psychophysical tasks (Rivest & Cavanaugh, 1996; Saarela & Landy, 2012).

**Figure 12:**
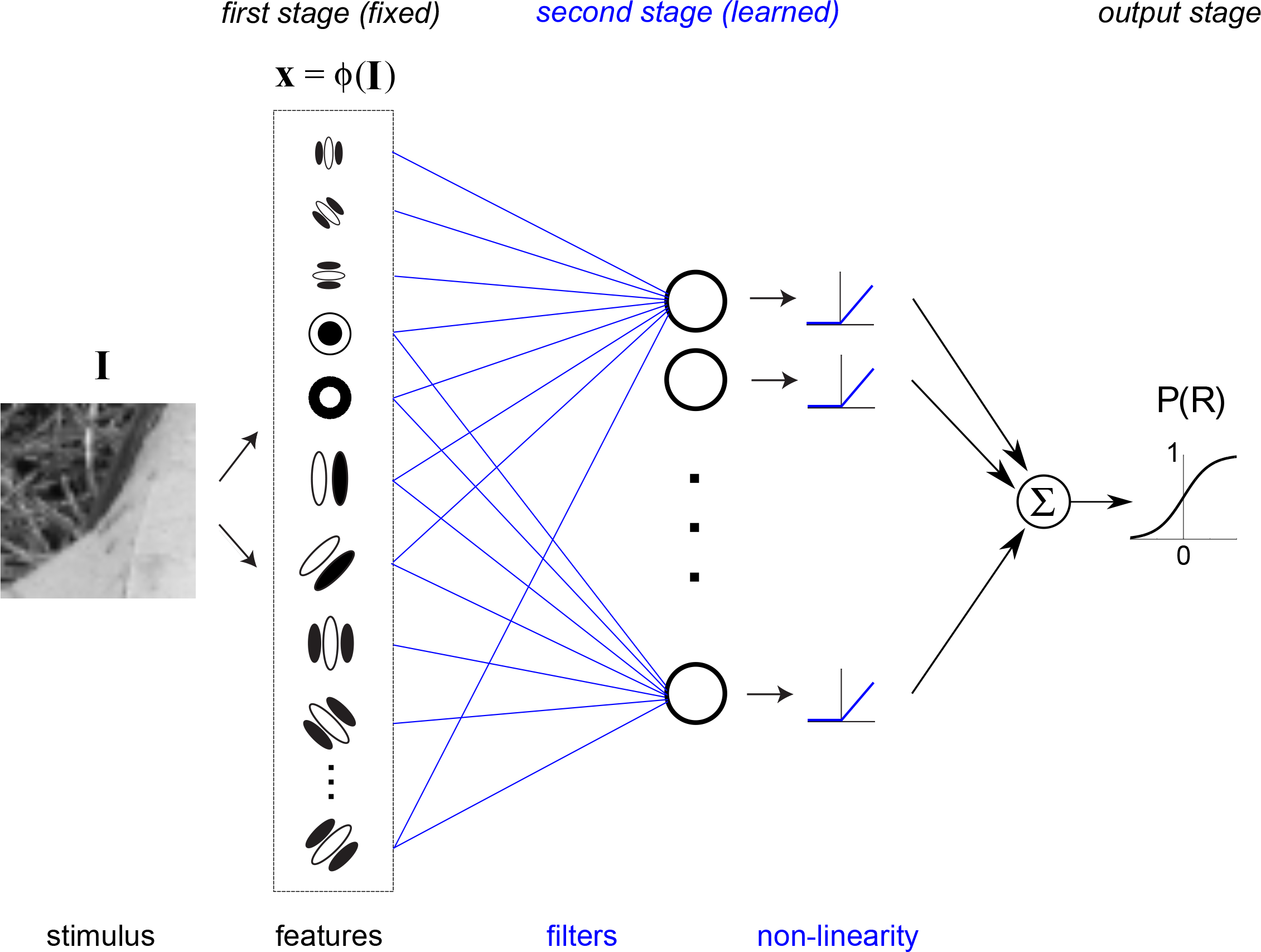
Hypothetical extension of our modeling framework with additional second-stage filters and a more complete implementation of the first-stage filters. Such a model could potentially be fit to human observer performance on boundary perception tasks involving complex stimuli such as natural occlusion boundaries.

An additional limitation of our model (and many other FRF variants) is the assumption of only a single stage of higher-order filtering beyond the first stage. Recent neurophysiological and fMRI studies have suggested that texture selectivity may develop progressively at successive stages of the visual hierarchy from V1 to V2 to V4 (Freeman et al. 2013; Goda et. al. 2014; Okazawa, Tajima, & Komatsu, 2015, 2016; Ziemba et al. 2016; Kohler et al. 2016). In studies using convolutional neural networks to solve image recognition tasks, texture-sensitive units often develop gradually over several processing layers (Yamins et al. 2014; Kahligh-Razavi & Kriegeskorte, 2014; Zeiler & Fergus, 2014; Yamins & DiCarlo, 2016). Furthermore, computer vision work on boundary detection has also found it useful to employ additional “texton” layers of processing (Martin et al., 2004). Therefore, it may be of interest for future work to consider models having additional intermediate layers of processing.

A number of previous studies have considered how information from an array of first-order visual filters may be pooled for perceptual decisions (e.g. Baker & Meese, 2014; Morgenstern & Elder 2012). In our analysis, we considered three different pooling rules inspired by previous work in computational vision science. We found that the spatial processing for the second-order vision tasks we examined was highly robust to choice of pooling rule (Supplementary Fig. S9), and that we could model our data accurately with different choices of pooling rule (Fig 7, Supplementary Figs. S12, S13). Therefore, we remain agnostic about the “best” model of pooling between the first and second stage filters, and suggest that this may be an interesting topic for future investigation.

Nevertheless, despite these limitations and our numerous modeling simplifications, in practice we found that for the tasks and the stimuli used here our simple FRF model (Fig. 2) was able to capture human performance quite well (Figs. 6-9).

### Comparison with previous psychophysical modeling

Despite the simplifications made in the present study, it represents a substantial methodological advance over previous efforts to model second-order visual perception. Although previous studies have developed image-computable FRF models, they have generally fixed not only the model architecture but also the shapes of the second-stage filters. The present study demonstrates how the second-stage filter shapes can be “learned” directly from experimental data in order to reveal the nature of spatial integration used in different tasks (Experiments 1 and 2).

Most previous FRF models in human psychophysics studies were conceptual or qualitative, without quantitative fitting of model parameters to data. Zavitz & Baker (2013, 2014) estimated free parameters of an FRF model by fitting threshold data as functions of stimulus parameters. Since thresholds are typically determined at only a few levels of a one-dimensional independent variable, this yields relatively few data points, greatly limiting the number of model parameters one can estimate. In our work, we directly estimate hundreds of model parameters by fitting stimulus-response data from thousands of psychophysical trials, enabling us to characterize the shapes of the second-stage filters.

In addition to extending the FRF modeling literature, our work also complements and extends previous classification image studies, which have generally been applied in the context of first-order vision (Neri & Levi, 2006; Murray, 2011). Although classification images yield image-computable predictive models of observer responses, relatively few studies validate these models by testing their predictive performance on a set of stimuli not used for model estimation. We tested model performance on novel stimuli not used for model estimation using both standard measures, e.g. percentage correct, as well as more advanced methods such as double-pass analysis (Neri, 2010; Hasan, Joosten & Neri 2012) and decision variable correlation (Sebastian & Geisler, 2018).

Finally, in contrast to most classification image studies, here we systematically address the issue of model comparison. Psychophysical system identification can be formulated in the framework of estimating a generalized linear model (Knobloch & Maloney, 2008a,b; Mineualt et al., 2009), which is unique up to choice of link function. Therefore, issues of model comparison (beyond coefficient shrinkage) do not naturally arise in this context. However, the FRF models relevant to second-order vision can vary greatly in architectural details (e.g., number of filtering layers, nonlinearities, connectivity, etc.), and model comparison is of great interest. In Experiment 3, we compare two competing FRF architectures making different assumptions about how the second-stage filters integrate across orientation channels, finding a preference for a model with early summation of first-order channels by the second-stage filters, consistent with previous psychophysical work (Kingdom, Prins & Hayes, 2003; Prins & Kingdom, 2002; Motoyoshi & Nishida, 2004).

### Edge-versus region-based segmentation

We applied our methodology to the question of whether or not observers take a region-based or an edge-based approach to segmenting a contrast boundary (Experiment 1). To the best of our knowledge, our experiment is the first time the issue of edge/region-based processing has been addressed for boundaries defined by differences in contrast, as it has for other second-order boundaries (Nothdurft, 1985; Gurnsey & Laundy 1992; Wolfson & Landy, 1998).

This question is of particular interest in light of several studies that have revealed possible neurophysiological mechanisms for contrast-boundary detection (Tanaka & Ohzawa, 2006; Rosenberg & Issa, 2010; Li et al. 2014; Gharat & Baker, 2016). This work has demonstrated a population of contrast-modulation (CM) tuned neurons in early visual cortex, which exhibit band-pass spatial frequency and orientation tuning to second-order modulation envelope frequency, consistent with a Gabor-shaped second-stage filter applied to the outputs of a bank of first-stage filters selectively tuned to the carrier pattern (Baker, 1999).

If such CM-responsive neurons were the mechanism responsible for psychophysical detection of CM boundaries, one might expect the 1-D weighting profiles to have a Gabor-like shape, placing greater weight on regions near the boundary. However, we find that this was not the case and that all areas of the image are weighted equally for the orientation-identification task (Experiment 1, Figs. 3, 4). Furthermore, in an additional control manipulation where we increased the carrier frequency in order to make sure CM neurons with smaller RFs would be relatively more engaged (Experiment 1-HFC), we still did not see any change in the type of spatial integration (Fig. 5). These findings are consistent with at least three distinct possibilities (not mutually exclusive) for the underlying neural mechanisms. One possibility is that task performance reflects CM-responsive neurons, like those described in V2 with large receptive fields (> 4 dva) encompassing the entire stimulus used in our experiments. Such neurons would prefer a carrier frequency much higher than their envelope frequency, which is plausible since primate V2 CM-responsive neurons can have preferred carrier:envelope ratios of up to 40:1 (Li et al., 2014). A second possibility is that psychophysical performance is mediated by a range of Gabor-like second-order mechanisms covering a range of spatial scales, and that our measured 1-d profiles reflect an ensemble of their contributions. Another possibility is that an important role is played by other neurons sensitive to texture patterns, such as those described in ventral stream areas (Hanazawa & Komatsu, 2001; Okazawa et al., 2015, 2016; Ziemba et al., 2016).

Clearly, the fact that we can readily distinguish non-juxtaposed textures implies that there must be mechanisms for texture representation that operate in the absence of a boundary. However, in some cases texture segmentation can benefit from boundary information, as in the textural Craik-Cornsweet illusion (Nothdurft, 1985). For example, Wolfson & Landy (1998) demonstrated that direct juxtaposition of textures having different orientations gives rise to local junction or corner cues that can support segmentation. Such enhanced segmentation might be mediated by neurons such as those described by Hegde & Van Essen (2000) and Anzai, Peng & Van Essen (2007), which are selective for such cues.

However, other work on texture segmentation has shown a minimal contribution of information along the boundary. A study using natural image patches in a task of discriminating occlusion boundaries from uniform textures found that removing the boundary region had little effect on performance (DiMattina et al., 2012). However, performance on natural occlusion edge and junction detection tasks can be seriously degraded when textural information, which is by its nature defined over a region, is removed (McDermott 2004; DiMattina et al., 2012). Similarly, segmentation of complex textures defined by letters shows little effect of boundary information (Gurnsey & Laundry, 1992).

In general, the answer to this question is most likely dependent on the particular stimulus and the nature of the psychophysical task, as indicated by the very different results of our first two Experiments. In our view, the question of “edge” versus “region”-based processing should be treated with some caution, since clearly for a very large stimulus extending into the far periphery there will be reduced weighting at the largest eccentricities, since visual acuity falls off sharply outside the fovea. In such a case the real question of interest would be the shape of the fall-off in weight as a function of stimulus eccentricity, and how this fall-off may relate to putative neural codes. Our method is well suited to address such issues for many different kinds of second-order boundaries and tasks, much as standard classification images can be similarly used for first-order boundaries.

In Experiment 1 every part of the image is useful for performing the task, and hence the ideal observer will integrate information from all image regions. In Experiment 2, the only useful information for performing the task is near the boundary and so an ideal observer will be indifferent to the stimulus elsewhere. In terms of possible mechanisms, we might expect CM neurons with larger receptive fields to be engaged in Experiment 1, whereas the most useful neural substrate in Experiment 2 might be CM-responsive neurons with smaller receptive fields. This is because CM neurons with larger vs. smaller receptive fields would be the ones whose responses best distinguish the two stimulus categories in Experiment 1 vs. 2, respectively.

### Integration of heterogeneous first-stage channels

A common ingredient to the various FRF models proposed to account for psychophysical texture segmentation data is a set of fine-scale first-stage filters summed by one or more second-stage filters defined on a much larger spatial scale. However one aspect in which such models differ is whether the individual second-stage filters analyze first-stage filter responses at a single orientation/spatial frequency, or whether they integrate across heterogeneous first-stage channels. In one standard model of FRF processing, each second-stage filter only analyzes the outputs of a single first-stage orientation/spatial frequency channel (Malik & Perona, 1990; Wilson, 1993; Landy & Oruc, 2002). However a number of psychophysical results have called this model into question. One study using a sub-threshold summation paradigm found support for a model utilizing linear integration across carrier orientation for detecting contrast modulations (Motoyoshi & Nishida, 2004). Another study making use of energy-frequency analysis revealed the existence of orientation-opponent mechanisms in second-order vision (Motoyoshi & Kingdom, 2003). Other work has suggested at least partly independent mechanisms for detecting second-order modulations defined by contrast and spatial/frequency orientation, with the suggestion that contrast modulations may be pooled across first-stage channels (Kingdom, Prins & Hayes 2003). In Experiment 3 we find support for a model with second-stage filters that integrate across first-stage orientation channels, consistent with the latter psychophysical studies. Although CM-responsive neurons described so far exhibit narrow tuning for carrier orientation (Li et al., 2014), this discrepancy could be due to the use of grating carriers rather than multi-orientation micropatterns. This suggests that that possible effects of carrier orientation bandwidth might be worth future investigation, both in human psychophysics and for CM-responsive neurons.

## Conclusions

Statistical machine learning has in recent years become a powerful and widely applied methodology in vision and in computational neuroscience. Our work demonstrates that the same machine learning methodology used to characterize neural coding (Wu et al., 2006; Willmore et al., 2010; Kriegskorte, 2015; Yamins & DiCarlo, 2016; Antolik et al., 2016) and first-order visual processing (Mineault et al. 2009; Murray, 2011) can be fruitfully applied to extend psychophysical system identification approaches to complex visual stimuli, including second-order boundaries. We anticipate that future applications of this approach may lead to a better understanding of how multiple first- and second-order cues are combined by human observers to detect natural boundaries, and provide insights into neural and computational mechanisms of image region segmentation.

## Methods

### Experimental Paradigms

#### Stimuli and psychophysical task

Contrast-modulated texture boundary stimuli were created similarly to those of Zavitz & Baker (2013), by multiplying a large (predominantly low spatial frequency) contrast modulation envelope by a high spatial frequency carrier (texture) pattern (Fig. 1a, b). The carrier patterns were produced by adding odd-phase Gabor micropatterns at random locations in an over-size larger image, which was subsequently cropped to the desired size. Only texture patterns having sufficient uniformity in root-mean-squared (rms) contrast were utilized, following the uniformity criterion of Zavitz & Baker (2013). To produce contrast-modulated stimuli, carrier textures were modulated by a half-disc oriented either right oblique (clockwise) or left-oblique (counter-clockwise), and then multiplied by a circular raised cosine window. The rms contrast of the carrier patterns was set to 14%, which is 6 dB above previously observed average carrier detection thresholds (Arsenault, Yonessi & Baker, 2011). Examples of right-oblique boundaries with varying carrier mircopattern densities and depths of contrast modulation are shown in Fig. 1b. Boundaries between micropattern textures like these have been used in previous psychophysics studies (Zavitz & Baker, 2013, 2014), and offer flexibility to produce a wide variety of naturalistic texture patterns amenable to parametric variation.

In all experiments, observers performed a single-interval 2AFC task to determine whether a briefly displayed (100 msec) boundary stimulus was contrast-modulated in a right- or left-oblique orientation. Envelope phase was randomized across trials, to preclude its use as a cue. Visual feedback was provided after each trial to maximize observer performance. Data was collected over multiple (4-5) sessions of 30-60 minutes, conducted over several days. An initial training session was dedicated to learning the orientation identification task (+/- 45 degree) by performing a short version (300 trials) at various micropattern densities (256, 1024, 2048 patterns or 512, 1024, 2048 patterns). Consistent with previous studies on second-order edge detection (Zavitz & Baker, 2013, 2014), we found in preliminary experiments that task performance was best for carrier texture patterns having a high density of micropatterns. Therefore we used relatively high micropattern densities (2048 per 256×256 image for Experiments 1, 2 and 512 for Experiment 3), higher than in previous studies (Zavitz & Baker, 2013), with the goal of optimizing observer performance.

#### Psychophysical observers

Observers were author CJD and 7 members of his research group (adult undergraduate students) who were initially naïve to the experimental purposes. For Experiment 1, 4 additional observers were recruited from two Psychology classes (EXP-3202, PSB-4002) at Florida Gulf Coast University (FGCU). All observers had normal or corrected-to-normal acuity. All observers provided written informed consent and all procedures were approved beforehand by the IRBs at FGCU (Protocol 2014-01) and McGill University (Protocol MED-B-96-279), in accordance with the Declaration of Helsinki.

#### Displays and experimental control

FGCU experiments were run in one of two windowless vision psychophysics chambers with no light sources other than the computer. Chamber 1 contained a factory-calibrated (gamma-corrected) LCD (Cambridge Research Systems, Display++, 120Hz, 1920×1080 pixels, 100 cd/m^2^). Chamber 2 contained a gamma-corrected CRT (SONY FD-500 Trinitron, 75 Hz, 1024×768 pixels, 59.2 cd/m^2^) driven by a Bits# stimulus processor (Cambridge Research Systems). In each chamber the display was controlled by a Dell OptiPlex 9020 with an NVIDIA GeForce GTX 645 graphics card. Observers viewed the stimuli binocularly, with a chin rest, at a distance of 133 cm, so that the 256×256 stimulus image subtended approximately 4 deg. visual angle (dva). McGill data was collected from author CJD for Experiments 2 and 3. Stimuli were displayed on a CRT (SONY Multiscan G400, 75 Hz, 1024×768 pixels, 53 cd/m^2^) with gamma correction through the color lookup tables. This monitor was driven by an Apple Mac Pro 4,4 (2.26 GHz Quad Core) hosting an NVIDIA GeForce GT 120 graphics card. Binocular viewing was at a distance of 125 cm, so that the stimulus subtended 4 deg, and experiments were conducted in a room with normal lighting. We observed remarkable consistency in our results from the three setups, despite differences in monitor characteristics and room lighting conditions, so all data was pooled for analysis. Experiments were controlled using custom-authored software written in the MATLAB^®^ programming language (MathWorks^®^, Inc.), making use of routines from the PsychToolbox^®^ Version 3.0.12 (Brainard, 1997; Pelli, 1997; Kleiner et al., 2007). To speed up experiments, carrier patterns were pre-computed and stored at 8 bit precision on the host computer. A unique set of pre-computed carrier patterns was used for each experiment, so that a given observer never saw the same carrier pattern more than once.

#### Experiment 1: Orientation identification

The stimuli for Experiment 1 were contrast-modulated discs (Fig. 1b). Stimuli were 256 × 256 pixels in size with a carrier pattern comprised of 2048 randomly placed, vertically oriented (0 degrees) Gabor micropattern stimuli (32 pixels or 0.5 dva., 1:1 aspect ratio). This carrier was modulated by an obliquely oriented half-disc, graduated by a cosine taper to avoid introduction of first-order artifacts. In this task, the two stimulus alternatives had right/left-oblique (+/- 45 deg.) boundaries.

In the main version of Experiment 1, for 7 observers, contrast modulation depths were varied across 9 logarithmically spaced levels from 8% to 32%, centered at 16%, plus 0% modulation (Experiment 1-VAR), using a method of constant stimuli. This set of levels resulted in trials at or near threshold for most observers, while still including some relatively easy trials. After an initial training session, observers each performed a total of 4000 trials (4 blocks of 1000 trials, 100 trials per level) over the course of multiple sessions spaced over several days. For one observer (CJD), 2975 micropatterns were used instead of 2048.

For 3 observers (CJD and naïve observers MAK and JJF), contrast modulation depths were fixed at a value slightly above each observer’s measured threshold (Experiment 1-FIX), with the goal of obtaining near-threshold task performance of approximately 80% correct (Supplementary Fig. S12). This is more typical of most classification image literature (Murray, 2011), which often fixes stimulus strengths at or near task performance thresholds. These observers performed 6000 trials (6 blocks of 1000 trials) over multiple sessions spaced across several days.

As an additional variant of Experiment 1, we used a carrier texture composed of micropatterns with a much higher spatial frequency (8 pixels or 0.125 dva.) and a higher density (8192), with rms contrast remaining at 14%. Experiment 1-HFC was run on three naïve observers (LJW, JMM, VHB) tested at variable modulation contrasts (same levels as Experiment 1-VAR).

#### Experiment 2: Fine orientation discrimination

In order to show the general applicability of our methodology to tasks where observers may employ an edge-based spatial summation, we ran a fine orientation discrimination task in which observers determined whether a clearly visible contrast-modulated boundary (same carrier parameters as in Experiment 1) was tilted slightly left-vs right-oblique relative to vertical. Experiment 2 is complementary to Experiment 1, in which the orientation difference is very large (+/- 45 degrees from vertical), while the contrast modulation is near detection threshold, whereas in Experiment 2 the contrast modulation is well above detection threshold, but the orientation difference is very small. This experiment was run on 2 observers, one of whom (JJF) was naïve to the purpose of the experiment. In a preliminary experiment at 32% contrast modulation, a method of constant stimuli was used to measure a just-noticeable-difference (JND) for orientation discrimination (approximately 75% performance) for each observer (JJF: 6 deg., CJD: 7 deg.). Then in subsequent sessions, observers performed the fine orientation discrimination at this value for a set of stimuli whose contrast modulation was varied from 16% to 64% (center = 32%) with a method of constant stimuli. Observers performed 6000 trials of the task over multiple sessions spaced over multiple days (6 blocks of 1000 trials, 100 trials per level). We refer to this experiment as Experiment 2-VAR. For two naïve observers (VHB, JJF) we also ran Experiment 2-FIX, where the orientation difference was fixed at each observer’s JND.

#### Experiment 3: Orientation identification with a more complex carrier texture

Experiment 3 was the same as Experiment 1-VAR except using a carrier texture containing micropatterns with two orthogonal orientations, as in Fig. 1a (bottom). For this experiment the micropatterns were elongated (2:1 aspect ratio instead of 1:1 as in Experiments 1, 2), to give a narrower orientation bandwidth. In addition the density was reduced (512 micropatterns instead of 2048), to create a stronger percept of oriented structure in the carrier texture.

### Modeling approach

#### Modeling framework

Most previous psychophysical SI methods assume that the decision variable arises from a weighted linear sum of the values (e.g. pixel luminances) of the sensory stimulus (Ahumada, 2002; Murray, 2011), essentially modeling the entire chain of brain mechanisms involved in stimulus-processing and decision-making by a single linear filter followed by a threshold. Here we propose a more general approach which we refer to as *psychophysical network modeling*. In this scheme, the decision variable is a function of the stimulus-dependent activities of a set of "hidden units", each of which may be sensitive to different aspects of the stimulus. This idea closely parallels recent modeling of neural responses by combining the outputs of multiple nonlinear subunits selective for different stimulus features (Willmore, Prenger & Gallant, 2010; Vintch, Simoncelli & Movshon, 2015; Harper et al. 2016; Rowenkamp & Sharpee, 2017).

Given a classification task for a stimulus **x** ∈ ℝ^*d*^ which elicits one of two behavioral alternatives (^*b*^ = 1, ^*b*^ = −1), one may define a generic psychophysical model as

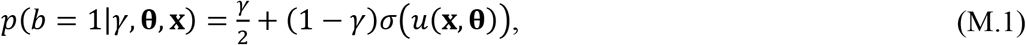

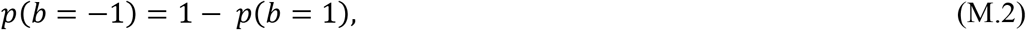

where *σ* is a psychometric function, *u*(**x**, **θ**) denotes the stimulus-dependent input to the psychometric function whose parameters **θ** characterize the sensory processing model, and *γ* denotes the lapse rate. The stimulus inputs **x** may be either the stimulus parameterization itself (e.g. pixel luminances), **I**, or a transformed version of the stimulus parameterization (**x** = *ϕ*(**I**)). We define a *psychophysical network model* by making the additional assumption that the input *u*(**x**, **θ**) to the psychometric function is some function of the activities *s*_1_, …, *s*_*K*_ of a set of *K* hidden units, i.e.

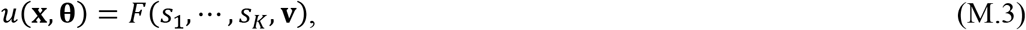

where **v** is a set of parameters characterizing how the *s*_*i*_ are combined. Perhaps the simplest combination rule is a weighted linear summation, i.e.

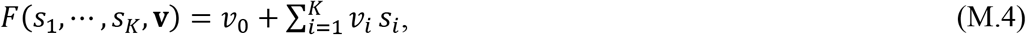

Note that we can have more than 2 hidden units (K > 2) even though there are only two behavioral responses (e.g. model of Fig.10a). We assume that the hidden unit activities are a nonlinear function of the projection of the features **x** onto some filter **w**_*i*_,

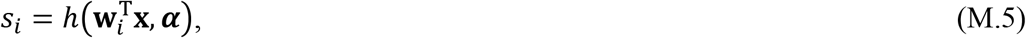

where **α**parameterizes the shape of the nonlinearity. Combining Eq. (M.1), (M.4) and (M.5) yields a final expression for a psychophysical network model with linear cue combination

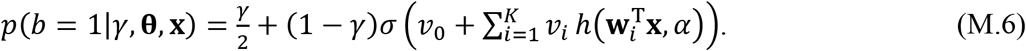

Various standard models used in machine learning are special cases of (M.6). For instance, standard binary logistic regression (Bishop, 2006) used for simple yes/no detection corresponds to the case of *γ* = 0 and a single hidden unit whose nonlinearity is a simple linear pass-through (*h*(*u*) = *u*) with fixed output weight *v*_1_ = 1, so that (M.6) reduces to

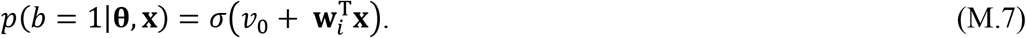

Likewise, with *γ* = 0 and the sigmoidal hidden unit nonlinearity *h*(*u*) = 1/(1 + *e*^−*u*^), this model (M.6) becomes a three-layer connectionist neural network (Rumelhart, Hinton, & Williams, 1986; Bishop, 2006).

#### Filter-rectify-filter model

A psychophysical network model for detecting whether a second-order edge is oriented left- or right-oblique is illustrated in Fig. 2. In this model, the inputs **x** are the down-sampled outputs of a bank of first-stage energy filters resembling V1 complex cells applied to the stimulus image **I**, a transformation which we represent as **x** = *ϕ*(**I**). There are two second-stage filters which serve as inputs to the hidden units. One hidden unit (L) learns a set of filter weights (blue lines) to detect a left-oblique edges from the pattern of activity in the first-stage subunits, while the other hidden unit (R) learns a set of filter weights to detect a right-oblique edge. The filter outputs are passed through a rectified power-law nonlinearity *h*(*u*) = |*u*|^*α*^ (blue curve) to generate the hidden unit activities *s*_*L*_, *s*_*R*_. Note it is important for *h*(*u*) to be even-symmetric, i.e. *h*(*u*) = *h*(−*u*), to provide envelope phase-invariance - otherwise the model would require four rather than two hidden units, since in our psychophysical experiment the envelope phase is randomized on each trial.

The hidden unit activities *s*_*L*_, *s*_*R*_ are combined, assuming fixed output weights (*v*_*R*_ = 1, *v*_*L*_ = −1), and a bias term *v*_0_ is added to obtain a decision variable *u* = *s*_*R*_ − *s*_*L*_ + *v*_0_. The fixed output weights are a consequence of our choice of a power-law nonlinearity, since for all inputs **x**, filters **w** and output weights *v*, the term *v*|**w**^T^**x**|^*α*^ is identical for all *v*, **w** satisfying *v*‖**w**‖^*α*^ = *C*, where *C* is some constant and ‖**w**‖ denotes the norm of **w** - see Ran & Hu (2017) and DiMattina & Zhang (2010). Many variants on this basic FRF model architecture are possible, and fitting multiple models to a given dataset could permit us to investigate questions about FRF model architecture (Motoyoshi & Nishida, 2004; Motoyoshi & Kingdom, 2007; Westrick & Landy, 2013).

A model of the form (M.6) arises naturally where there are two signals representing the two stimulus alternatives whose activities are compared to make a decision, for instance, the outputs of the two subunits in Fig. 2. Subtracting the activities *s*_1_ − *s*_2_ and adding a bias *v*_0_ and Gaussian internal noise *ε*~*N*(0, *ρ*^2^) models the proportion of times behavioral alternative *b* = 1 is chosen as

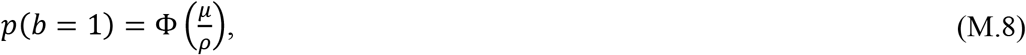

where Φ is the cumulative unit normal distribution and *μ* = *s*_1_ − *s*_2_ + *v*_0_. Using the approximation Φ(*u*) ≈ *σ*(*ku*), where *σ* is the sigmoid function and *k* is optimized to best approximate Φ(*u*), we can re-write (M.8) as

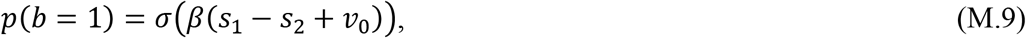

where 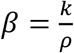 is inversely proportional to the internal noise *ρ*. Since 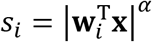, this inverse internal noise parameter *β* is simply absorbed into the magnitudes of the second-stage filter weights **w**_*i*_ and bias *v*_0_. Therefore, larger filter norms will correspond to larger values of *β* and lower internal noise. In the analysis here, the norms of the **w**_*i*_ were not solely determined by the internal noise since they were also constrained by the Bayesian prior. However, we found that the model provided an excellent fit to our psychophysical data without explicitly optimizing an additional internal noise parameter, so with the exception of only one analysis for Experiment 1-FIX (Supplementary Fig. S13) we did not optimize a noise parameter.

#### Model estimation and regularization

Here we simplify notation by combining all model parameters and the lapse rate into the single parameter vector **η**. Given psychophysical stimulus-response data 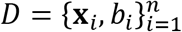, and prior constraints *p*_0_(**η**) (possibly uninformative) on the model parameters, we pose the estimation as a machine learning problem of finding the maximum a posteriori (MAP) estimate of **η**, given by:

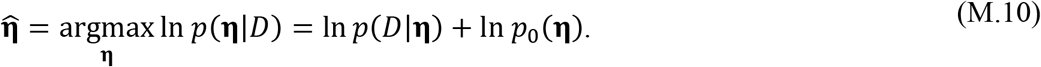

In this framework, the prior constraints *p*_0_(**η**) are often used to prevent over-fitting (Bishop, 2006), which can be a serious problem due to the relatively large number of parameters (hundreds or thousands) being estimated from a limited number (generally < 10K) of psychophysical trials. Various techniques have been applied to help reduce the problems of noise and over-fitting, including Bayesian priors (Mineault, Barthelme & Pack, 2009), radial averaging (Abbey & Eckstein, 2002) or using simplified low-dimensional stimuli (Li, Levi & Klein, 2004). Here we considered two different Bayesian methods for regularization of the second-stage filter weights. Under a prior assumption that the weights are from a multivariate Gaussian distribution, a “ridge-regression” penalty (Bishop, 2006) is applied to each of the second-stage filter weight vectors **w**_*i*_ (Fig. 2, blue lines) - this encourages small weight magnitudes, to prevent fitting noise. The prior is given by

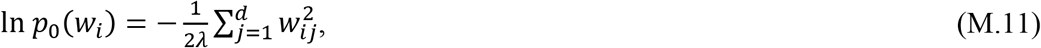

where *w*_*ij*_ denotes the *j*-th element of filter **w**_*i*_ and the hyperparameter *λ* controls the strength of the penalty (smaller *λ* induces a stronger penalty). Optimizing *λ* using *k*-fold cross-validation (Bishop, 2006), *λ* = 0.1 − 1.0 provided a good trade-off between fitting to the training data and generalizing to novel data in the test set in Experiment 1 - see Supplementary Figs. S1, S2. This parameter was re-optimized for each of the two models in Experiment 3 (which differed in architecture from those used in Experiments 1, 2) using the same procedure (Supplementary Fig. S19).

We also considered the effects of an additional prior *q*_0_(**w**_*i*_) favoring second-stage weights that form a spatially smooth filter (Wilmore & Smyth, 2003). Such a prior was implemented by constructing a matrix **S**_2_ whose rows are the 2-D discrete Laplace operator

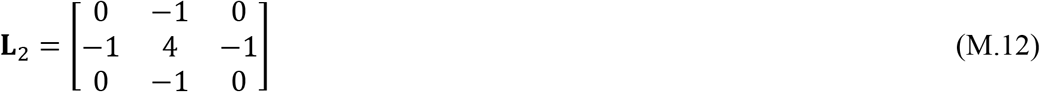

centered at every point in a *p* × *p* matrix of zeros and unwrapped into a vector of length ^*d*^ = *p*^2^, where *p* is the down-sampled image dimensionality. This smoothness prior is specified by

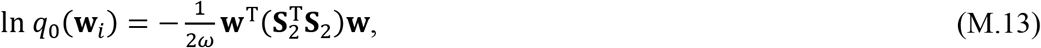

with hyper-parameter *ω* controlling the extent to which roughness is penalized (smaller *ω* induces a stronger penality). Optimizing *ω* using *k*-fold cross-validation with *λ* = 1, there was a slight improvement in generalization when the smoothness prior (M.13) was also included, with the value of *ω* =1 providing a good tradeoff between fitting the data and generalizing to novel test data in Experiment 1 (Supplementary Figs. S3, S4). Adding this additional smoothness constraint reduced fitting to the training dataset, but improved generalization of the model to novel data (Supplementary Figs. S5, S6).

#### Numerical optimization

Second-stage filter weights **w**_*i*_ and the exponent α of the hidden unit nonlinearity were estimated using alternating coordinate ascent in a manner similar to the EM algorithm (Dempster, Laird and Rubin, 1970). Weights were optimized for one filter **w**_*i*_ at a time while all other filters **w**_*j*≠*i*_ and the power law exponent α were fixed. The filter weights, starting with randomized initial values, were optimized using a quasi-Newton method (L-BFGS). To estimate the power law exponent α, all of the second-stage filters were fixed and a 1-D line search was performed over the range [0.1,4.0]. The optimizations for **w**i and α are not log-concave, which means that local maxima may be possible (Paninski, Pillow & Lewi, 2007). In practice we usually found robust and consistent estimates of model parameters across optimization runs starting from several starting points. We found that sometimes the search converged on a “local maximum” consisting of only a single non-zero filter. Therefore, as an additional precaution, after the first round of optimization (or at least 4 total rounds depending on the experiment) we always re-initialized the search at a point in parameter space where the filter that was smaller in magnitude was set to be mirror-symmetric to the current value of the larger one. The optimization was then allowed to proceed as usual for several more rounds before termination. This procedure had the effect of eliminating any problems with local minima.

In preliminary investigations the estimated lapse rate *γ* was always zero, so this parameter was not included in our final model. Previous work has shown the best estimates of lapse rates when a large number of easy trials are included (Prins, 2012). This was not the case in our study (which focused trials near threshold), nor in the general classification image literature, which tends to use near-threshold stimuli and does not typically estimate lapse rates (Murray, 2011).

## Appendix A

Here we derive a simple expression for the Bayes factor for two binomial models. We show using our formula that over large numbers of trials, a small difference in the accuracy of the two models can lead to a very strong preference for the more accurate model, as measured by the Bayes Factor.

Let *M*_1_, *M*_2_ be two binomial models, each predicting for a fixed stimulus, positive response (*r* = 1) probabilities of *p*_1_, *p*_2_ and negative response (*r* = 0) probabilities 1 − *p*_1_, 1 − *p*_2_. Assume that we perform *n* experimental trials and observe proportion *p*\* positive responses and (1 − *p*\*) negative responses. Let *D* = *r*_1_, …, *r*_*n*_ denote the observer responses. The Bayes Factor is

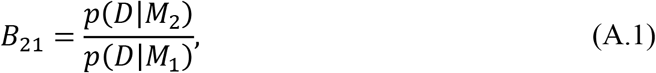

and taking the natural log we obtain

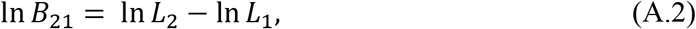

where *L*_*k*_ = *p*(*D*|*M*_*k*_) for (*k* = 1,2). Expanding the log-likelihoods, we obtain

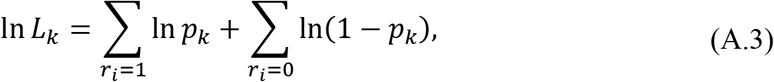

where *p*_*k*_ = *p*(*r* = 1|*M*_*k*_). Since there are *np*\* trials with positive responses and *n*(1 − *p*\*) trials with negative responses, we may simplify (A.3) to

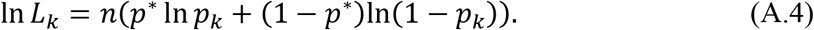

Noting that the bracketed term in (A.4) is simply the negative cross-entropy between two binomial models, and using the fact that the cross-entropy *C*(*p*, *q*) relates to the entropy *H*(*p*) and the KL divergence *D*_*KL*_(*p*‖*q*) by the relation *C*(*p*, *q*) = *H*(*p*) +*D*_*KL*_(*p*‖*q*), we can re-write (A.4) as

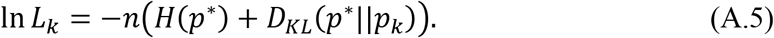

Plugging (A.5) into (A.2), the natural log of the Bayes factor becomes

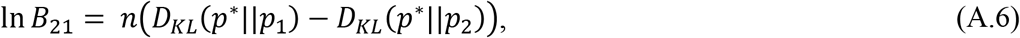

and we obtain the final expression

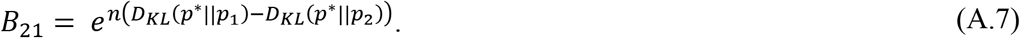

We see from the final result (A.7) that the Bayes factor magnitude is exponential in the number of trials (*n*), meaning that small differences in model performance (measured by the difference in the KL divergences) over a large number of trials can lead to very large values for the Bayes factor. Similarly, we see from (A.6) that the standard model selection criterion 2ln *B*_21_ (Kass and Raferty, 1995) grows linearly as *n*, meaning that a small difference in performance may accumulate over trials to produce large values of this criterion.

Concretely, consider the case where *p*\* = 0.75, *p*_1_ = 0.73, *p*_2_ = 0.74 for *n* = 5000 trials. Qualitatively, we see that both models are slightly inaccurate, with Model 2 agreeing with the observer more than Model 1, but with a difference of only 0.01. Applying formula (A.6) gives us ln *B*_21_ = 3.8458 and therefore 2ln *B*_21_ = 7.6916, corresponding to a “strong” preference for Model 2 according to standard criteria (Kass and Raferty, 1995). Re-working this example with a slightly larger difference in performance (0.02) by setting *p*_1_ = 0.72 yields a “very strong” preference for Model 2 (2ln *B*_21_ = 20.2224), and a 2.46 × 104 fold preference for Model 2. This effect is even more pronounced for predictions of very high (or low) success rates: setting *p*\* = 0.95, *p*_1_ = 0.92, *p*_2_ = 0.94 yields 2ln *B*_21_ = 60.4679, greatly exceeding the standard criterion (10) for “very strong” evidence in favor of Model 2.

## Acknowledgements

We thank the FGCU undergraduates in the Computational Perception Lab for help with data collection, Wilson S. Geisler for code implementing decision-variable correlation, Elizabeth Zavitz for micropattern texture stimuli, and Fred Kingdom for helpful discussions and comments. Supported by Canadian NSERC Grants OPG0001978 and RGPIN-2017-05292 to C.L.B. and a FGCU Professional Development Grant to C.D.

